# Alzheimer’s genetic risk factor *FERMT2* (Kindlin-2) controls axonal growth and synaptic plasticity in an APP-dependent manner

**DOI:** 10.1101/767194

**Authors:** Fanny Eysert, Audrey Coulon, Emmanuelle Boscher, Anaїs-Camille Vreulx, Amandine Flaig, Tiago Mendes, Sandrine Hughes, Benjamin Grenier-Boley, Xavier Hanoulle, Florie Demiautte, Charlotte Bauer, Mikael Marttinen, Mari Takalo, Philippe Amouyel, Shruti Desai, Ian Pike, Mikko Hiltunen, Frédéric Chécler, Mélissa Farinelli, Charlotte Delay, Nicolas Malmanche, Sébastien Hébert, Julie Dumont, Devrim Kilinc, Jean-Charles Lambert, Julien Chapuis

**Affiliations:** Univ. Lille, Inserm, CHU Lille, Institut Pasteur de Lille, U1167 - RID-AGE - Facteurs de risque et déterminants moléculaires des maladies liées au vieillissement, Lille 59019 France; Centre de recherche du CHU de Québec-Université Laval, CHUL, Axe Neurosciences, Québec, Canada; Faculté de médecine, Département de psychiatrie et de neurosciences, Université Laval, Québec, Canada; E-PHY-SCIENCE, Bioparc de Sophia Antipolis, 2400 route des Colles, Biot 06410 France; Univ. Lille, CNRS, UMR8576 - Labex DISTALZ, Villeneuve d’Ascq 59655 France; Université Côte d’Azur, Inserm, CNRS, IPMC, DistAlz Laboratory of Excellence, Valbonne, France; Institute of Biomedicine, University of Eastern Finland, Kuopio, Finland; Proteome Sciences plc, Hamilton House, London WC1H 9BB, United Kingdom

**Author notes:** Corresponding author: Julien Chapuis, PhD, Inserm UMR1167, Institut Pasteur de Lille, 1 rue du Pr. Calmette, Lille 59019 France. These authors share the last authorship (Jean-Charles Lambert, Julien Chapuis).

## Abstract

Although APP metabolism is being intensively investigated, a large fraction of its modulators are yet to be characterized. In this context, we combined two genome-wide high-content screenings to assess the functional impact of miRNAs and genes on APP metabolism and the signaling pathways involved. This approach highlighted the involvement of *FERMT2* (or Kindlin-2), a genetic risk factor of Alzheimer’s disease (AD), as a potential key modulator of axon guidance; a neuronal process that depends on the regulation of APP metabolism. We found that FERMT2 directly interacts with APP to modulate its metabolism and that FERMT2 under-expression impacts axonal growth, synaptic connectivity and long-term potentiation in an APP-dependent manner. Lastly, the rs7143400-T allele, which is associated with an increased AD risk and localized within the 3’UTR of FERMT2, induced a down-regulation of FERMT2 expression through binding of miR-4504 among others. This miRNA is mainly expressed in neurons and significantly overexpressed in AD brains compared to controls. Altogether, our data provide strong evidence for a detrimental effect of FERMT2 under-expression in neurons and insight on how this may influence AD pathogenesis.

## INTRODUCTION

AD is a neurodegenerative disease characterized by two main pathological hallmarks: (i) intracellular neurofibrillary tangles consisting of hyper-phosphorylated Tau proteins and (ii) extracellular amyloid plaques consisting of aggregates of β-amyloid (Aβ) peptides resulting from the processing of amyloid precursor protein (APP). Three main proteases (α-, β- and γ-secretases) are involved in APP processing through (i) the amyloidogenic pathway (β- and γ-secretases), leading to Aβ production, and (ii) the non-amyloidogenic pathway (α- and γ –secretases), which prevents Aβ generation by cleaving APP within the Aβ sequence (Checler 1995).

The identification of early-onset autosomal dominant AD-linked mutations in the genes for *APP* and presenilins (*PS1* and *PS2*, part of the γ-secretase), have placed abnormal APP metabolism at the center of the disease, further supporting the amyloid cascade hypothesis (Hardy 1997; Hardy & Selkoe 2002): the overproduction of Aβ peptides –especially the longer forms that are thought to be more neurotoxic– may lead to (or favor) Tau pathology and subsequent neuronal death.

Although the validity of the amyloid cascade hypothesis is debated (Morris et al. 2018), the importance of APP has recently been emphasized by the discovery of a rare *APP* mutation hampering Aβ production that lowers AD risk (Jonsson et al. 2012). Moreover, loss of functions variant in SORL1 (Sortilin-related receptor, L(DLR class A), which is a strong regulator of APP metabolism and Aβ production, is associated with early- and late-onset forms of AD (Nicolas et al. 2016; Andersen et al. 2005). Beyond Aβ production, the involvement of genetic risk factors such as APOE and TERM2 in modulation of Aβ aggregation and/or degradation/clearance has been proposed to be essential in the AD process (Kim et al. 2009; Jay et al. 2017). Recent high-throughput genomic approaches have also highlighted APP metabolism in the AD pathophysiological process: the main actors of APP metabolism, *e*.*g*., ADAM10 and APH1B (part of the γ-secretase complex), have been characterized as genetic determinants (Jansen et al. 2019; Kunkle et al. 2019), and numerous other genetic determinants have been described as potential modulators of APP metabolism (for a review, see (Dourlen et al. 2019)).

Among these genetic determinants, FERMT2 has been identified to be involved in APP metabolism using an agnostic, systematic approach, *i*.*e*., high content screening of 18,107 siRNA pools in HEK293 cells stably over-expressing an APP fusion protein (mCherry-APP^695WT^-YFP) that allows for the quantification of intracellular APP fragments (Chapuis et al. 2017). Following this initial screening, FERMT2 under-expression was then specifically associated with increasing levels of mature APP at the cell surface, where FERMT2 facilitates APP recycling, resulting in increased Aβ peptide production (Chapuis et al. 2017).

Little is known about FERMT2. This protein localizes to focal adhesions, where it is proposed to interact with β3 integrin and to be a major actor in integrin activation (Theodosiou et al. 2016). FERMT2 has been reported as a key protein involved in cardiac and skeletal muscle development (Dowling et al. 2008), and has been involved in cancer progression (Shen et al. 2012; Zhan et al. 2012; Sossey-Alaoui et al. 2019). However, despite of the fact that FERMT2 is a major genetic risk factor of AD, the physiological and/or pathophysiological roles of FERMT2 in the brain have not been identified. Within this background, we sought to determine how FERMT2 regulation impacts APP metabolism and/or AD risk and could be its involvement in neuronal functions.

## METHODS

### Cell culture

Human HeLa (RRID:CVCL_0030) and HEK293 (RRID:CVCL_0045) cells were respectively maintained in Eagle’s minimal essential medium (American Type Culture Collection, Teddington, UK) and DMEM/Ham’s F-12 1:1 medium (Life Technologies, Carlsbad, CA) supplemented with 10% heat-inactivated fetal bovine serum and 2 mM L-glutamine, penicilline (10 UI/mL)/Streptomycine (10 μg/mL).

### Microfluidic chip fabrication

Masters of multi-compartment microfluidic devices were fabricated through photolithography as previously described (Blasiak et al. 2015). Polydimethylosiloxane (PDMS; Sylgard 184; Dow Corning, Midland, MI) pads were replica molded (2 h at 70°C) and irreversibly bonded to glass coverslips *via* O_2_ plasma (Diener, Ebhausen, Germany). The devices were placed in plastic Petri dishes, wetted with dH_2_O, and UV sterilized for 30 min.

### Primary neuronal culture and viral transductions

Primary neuronal cultures were obtained from hippocampus or cortices of post-natal (P0) rats as described previously (Kaech & Banker 2006). Briefly, after the dissection of the brains, hippocampi were washed three times in HBSS (HBSS, 1 M HEPES, penicilline/streptomycine, 100 mM sodium pyruvate; Gibco) and were dissociated *via* trypsin digestion (2.5%, 37°C; Gibco) for 7 min. Next, hippocampi were incubated with DNAse (5 mg/mL; Sigma) for 1 min and washed again in MEM medium supplemented with 10% SVF, 1% Glutamax, 0.8% MEM vitamines, 0.5% penicilline/streptomycine and 0.45% D-glucose (Sigma). With a pipette, hippocampi were mechanically dissociated and resuspended in Neurobasal A medium (Gibco) supplemented with 2% B27 (Gibco) and 0.25% GlutaMax. 200,000 neurons were seeded per well in 24-well plates. 50,000 neurons were seeded in the somatic chamber of microfluidic devices, pre-coated with poly-L-lysine (0.1 mg/mL; Sigma) in borate buffer (0.31% boric acid, 0.475% sodium tetraborate, pH = 8.5). 0.1% EDTA (in H_2_O) was added to the Petri dishes containing microfluidic devices to minimize evaporation. The culture medium was refreshed every 3 d. Neurons were maintained at 37°C in a humidified 5% CO_2_ incubator.

### Lentiviral transductions

Lentiviral transductions were carried out at 1 day *in vitro* (DIV1) with a multiplicity of infection (MOI) of 10. In the case of co-transduction, MOI of 5 was used for each lentivirus. Briefly, lentiviruses were diluted in culture medium containing 4 μg/mL polybrene (hexadimethrine bromide; Sigma) and were added to the cells. After 6 h of transduction, lentivirus suspension was replaced with fresh medium. The following lentiviruses were used for transduction: Mission shRNA vectors (Sigma) shNT (Non-Mammalian shRNA Control, SHC002), shFERMT2 (TRCN0000191859), shAPP (TRCN0000006707) and pLenti6 empty vectors (Mock) or including human FERMT2^WT^ or FERMT2^QW^ cDNA sequences. LifeAct-Ruby lentivirus (pLenti.PGK.LifeAct-Ruby.W: RRID:Addgene_51009) was a kind gift from Rusty Lansford.

### RFLP genotyping

Genomic DNA in the vicinity of the rs7143400 was amplified by PCR using the following primers 5’-GGTTGGGTGTGAATAGGAAT-3’ and 5’-TGCATGCCTGATTTATTTGG-3’ before digestion with Tsp45I enzyme (Thermo Scientific). Finally, treated PCR products were analyzed in 2% agarose gel to visualize the cleavage bands.

### Designing CRISPR/Cas9 and genome editing

gRNA sequences were predicted by Benchling (http://www.benchling.com) and cloned into the GeneArt CRISPR OFP Nuclease Vector (ThermoFischer Scientific) allowing Cas9 and gRNA expression. Homology directed repair was induced by co-transfection of 71 pb double-strained DNA oligonucleotide template including rs7143400-T allele in HEK293 cells (**Supplementary Fig. 1**). HEK293 clones were isolated by limiting dilution before RFLP genotyping. Sequence integrity of the *FERMT2* 3’UTR and predicted potential off-target sites were validated by Sanger sequencing (**Supplementary Fig. 1**).

**Fig. 1.**
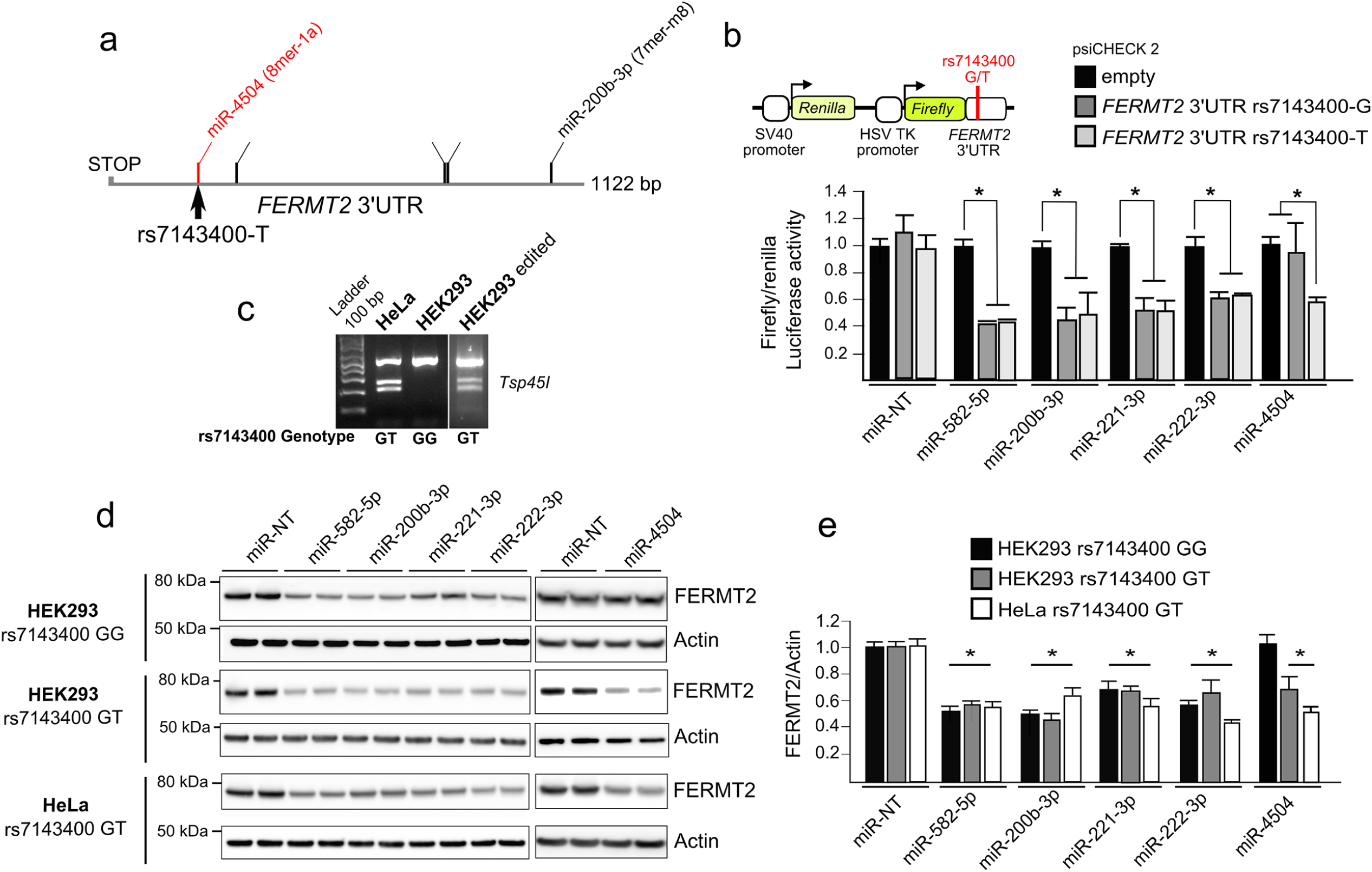
Validation of functional miRNAs targeting *FERMT2* 3’UTR. **a**. Relative positions of miRNA target sites on *FERMT2* 3’UTR. The target site created by the rs7143400-T allele, which is associated with AD risk, is shown in red. **b**. Luciferase activity of FERMT2 3’UTR carrying either the rs7143400-G or the rs7143400-T allele in HEK-293 cells co-transfected with a non-targeting miRNA (miR-NT) or 5 miRNA mimics. Data are expressed relative to the miR-NT **c**. RFLP genotyping of HeLa and HEK293 cell lines edited or not for the rs7143400 *via* CRISPR-Cas9 (**Supplementary Fig. 1**). **d**. Endogenous FERMT2 expression levels were assessed by Western blot using indicated cell extracts following transient transfection with a non-targeting miR (miR-NT) or with the indicated miR for 72 h. **e**. WB quantifications from three independent experiments as in ***d***. Data given in mean ± SD. * *p* < 0.05, non-parametric test compared to miR-NT condition.

### Visualization of miRNA expression at the single-cell level

To visualize RNA molecules by fluorescence at single-cell resolution and quantify gene expression, we used ViewRNA Cell Plus Assay kit (ThermoFischer Scientific) according to the manufacturer’s instructions. Briefly, after fixation and permeabilization, cells were washed 3× with PBS containing RNAse inhibitor and were incubated with probes directed against specific mRNA or miRNA for 2 h at 40°C. After washes, probes were amplified, first, in a pre-amplifier solution and second, in an amplifier solution, both for 1 h at 40°C. Then, cells were incubated with nucleotide probes stained with different fluorophores allowing the detection of mRNA or miRNA puncta. This approach was coupled with immunofluorescence experiments described.

### miRNA quantification in human brain samples

This study was approved by CHU de Québec – Université Laval Research Ethics Committee (#2017-3017). Frozen human brain tissue (0.5–1.2 g per sample) was obtained from the Harvard Brain Tissue Resource Center in Belmont, USA, the Brain Endowment Bank in Florida, USA, and the Human Brain and Spinal Fluid Resource Center in Los Angeles, USA, *via* NIH Neurobiobank. The cohort of patients included non-dementia controls (N = 30) and AD cases (N = 52) based on neuropathological diagnosis. Upon receipt of the specimens, frozen post-mortem parietal coxtex (BA39) was crushed using a biopulverizer prior to RNA extraction and analysis.

Total RNA was extracted from brain tissue (**Supplementary Table 1**) using TRIzol reagent (Ambion, 15596018) according to the manufacturer’s instructions. miRNA quantifications were done using the TaqMan miR Reverse Transcription Kit (Applied Biosystem, Burlington, Canada) and TaqMan Universal Master Mix (Applied Biosystem, 4324018) following the manufacturer’s instructions. Primers were purchased from ThermoFisher (miR-4504 ID: 464271_mat; RNU48 ID: 001006; miR-222-3p ID: 002276). MiR-4504 and miR-222 levels were normalized to RNU48. The relative amounts of each mature miRNA were calculated using the comparative Ct (2^-^ΔΔCt) method (Smith et al. 2011).

**Table 1.** Results of pathway enrichment analyses. **a**. The 10 most likely canonical pathways identified after pathway enrichment analysis of 41 miRNAs that strongly modulate APP metabolism using DIANA Tools mirPath (v3.0) **b**. The 10 most likely canonical pathways identified after pathway enrichment analysis of 132 genes targeted by 41 miRNAs (see Supplementary Methods for details)

| **KEGG pathway** | ***p*-value** | **#miRNAs** |
| --- | --- | --- |
| Axon guidance | 4.70E-08 | 48 |
| Proteoglycans in cancer | 1.82E-06 | 49 |
| Hippo signaling pathway | 4.10E-06 | 50 |
| Fatty acid biosynthesis | 8.10E-05 | 11 |
| Glutamatergic synapse | 8.56E-05 | 48 |
| GABAergic synapse | 1.10E-04 | 48 |
| AMPK signaling pathway | 2.20E-04 | 46 |
| TGF-beta signaling pathway | 2.50E-04 | 45 |
| Thyroid hormone signaling pathway | 3.90E-04 | 48 |
| Adrenergic signaling in cardiomyocytes | 5.00E-04 | 49 |

| **KEGG pathway** | ***p*-value** | **#genes** |
| --- | --- | --- |
| Axon guidance | 0.0014 | 19 |
| Ubiquitin mediated proteolysis | 0.010 | 16 |
| Circadian rhythm | 0.036 | 8 |

### Immunoblotting and Aβ quantification

Equal amounts (10-25 μg) of cell lysate were collected in RIPA buffer (1 M Tris, 1.5 M NaCl, 0.1% NP-40, 10% SDS, 100 mM sodium orthovanadate, 0.5% sodium deoxycholate, pH = 7.4) containing protease inhibitors (Complete mini; Roche Applied Science, Penzberg, Germany), lithium dodecyl sulfate (LDS), and reducing agent (Invitrogen). Samples were denaturated and analyzed using SDS-PAGE and the following antibodies: human FERMT2 (RRID:AB_10727911), amyloid precursor protein C-terminal domain (RRID:AB_258409), actin (RRID:AB_476692), β-amyloid clone 6E10 (RRID:AB_662798), β-amyloid clone 4G8 (RRID:AB_662812), Synaptophysin I (RRID:AB_887824), PSD95 (RRID:AB_2619800), GAPDH (RRID:AB_10615768). Extracellular culture media were collected in order to dose secreted Aβ using Alpha-LISA assays (AlphaLISA Amyloid β_1-X_ Kit, AL288C, PerkinElmer) according to the manufacturer’s instructions.

### Cell surface biotinylation

HEK293-APP^695WT^ cells were transfected with PCDNA4.1, FERMT2^WT^ or FERMT2^QW^ (PCDNA4/HisMax, ThermoScientific V86420) for 48 h. Next, cell surface proteins were biotinylated using sulfo-NHS-SS-biotine (sulfosuccinimidyl-20(biotinamido)ethyl-1,3-dithiopropionate) for 30 min at 4°C according to the manufacturer’s instructions (Cell Surface Protein Isolation Kit, Pierce, 89881). Then, cells were lysed and immunoprecipitated with streptavidin beads. Precipitated proteins were eluted from the beads with loading buffer containing 50 mM DTT, heated for 10 min at 95°C and analyzed by WB.

### Co-immunoprecipitation

Equal amounts of protein were collected in co-immunoprecipitation buffer (10 mM HEPES, 140 mM NaCl, 0.5% NP-40, pH = 7.4) containing protease inhibitors (Complete mini, Roche Applied Science) and phosphatase inhibitor (100 mM sodium orthovanadate) and incubated with the primary β-amyloid antibody clone 4G8 (RRID:AB_662812) overnight, with gentle rocking. Production of recombinant C100 fragment was performed as previously described (Sevalle et al. 2009). Co-immunoprecipitation was carried out using Pierce Protein A/G magnetic beads kit (Thermo Scientific, 88802) according to the manufacturer’s instructions. Samples with proteins and antibody complexes were incubated with 25 μL (0.25 mg) of A/G magnetic beads previously washed with co-immunoprecipitation buffer. After 1 h of incubation at 4°C, the magnetic beads were washed 3×, resuspended with loading buffer (LDS and reducing agent) for 10 min at RT, and analyzed by WB.

### Immunofluorescence and PLA

Cells were fixed in 4% paraformaldehyde (PFA) for 15 min, washed 3× with PBS, and permeabilized for 5 min with 0.3% Triton X-100. Cells were incubated with 5% normal donkey serum for 2 h at RT before overnight incubation with the following primary antibodies: human FERMT2 (RRID:AB_10727911), Kindlin2 (RRID:AB_2278298), amyloid precursor protein C-terminal domain (RRID:AB_258409), APP A4 clone 22C11 (RRID:AB_94882), Synaptophysin I (RRID:AB_887824), PSD95 (RRID:AB_2619800), Homer (RRID:AB_2631222), α-Tubulin (RRID:AB_2210391). The cells were then washed 3× with PBS and incubated with the following secondary antibodies raised in donkey (AlexaFluor-conjugated AffiniPure Fragment 405, 488, 594 or 647, Jackson ImmunoResearch), 1:10,000 Hoechst 33342, or 1/40 SiR-Actin probe (SC001, Spirochrome). Alternatively, Kindlin2 (RRID:AB_2278298) and APP A4 22C11 (RRID:AB_94882) antibodies were used for proximity ligation assay (PLA) according to manufacturer’s instructions (Duolink®, Olink Bioscience).

### Live-cell microscopy for axon elongation and actin dynamics

After DIV5, once the axons reached the axonal chamber of microfluidic devices, the culture medium was replaced with Neurobasal A without phenol red, supplemented with GlutaMax, 2% B_27_, and 25 mM HEPES. Phase-contrast images of growing axons were acquired every 10 min for 110 min using Zeiss AxioObserver Z1 microscope equipped with a Prime 95B Scientific CMOS (Photometrics, Tucson,AZ) camera and 32× objective. Movies were analyzed using Fiji MTrack J Plugin (Meijering et al. 2012) to determine the axon growth speed.

To visualize filamentous actin (F-actin) dynamics in the growth cones of elongating axons, neurons were co-transducted with LifeAct-Ruby at DIV1. At DIV5, growth cones expressing LifeAct-Ruby were imaged using a Nikon microscope equipped with Yokogawa spinning disk system and a Nikon CFI Apochromat 100× TIRF objective (NA 1.49), in live superresolution mode (66 nm/px). Processed movies were analyzed using Imaris (Bitplane, Zurich, Switzerland) surface tracking tool to obtain the speed and direction of F-actin puncta undergoing actin retrograde flow.

### Synaptosome extraction

To verify the presence of proteins at the synaptic level we did a subcellular fractionation as previously described (Frandemiche et al. 2014). Briefly, cortical neurons were resuspended in a solution (0.32 M sucrose and 10 mM HEPES, pH = 7.4) and were centrifuged at 1,000× *g* for 10 min to remove nuclei and debris. The supernatant was centrifuged at 12,000× *g* for 20 min to remove the cytosolic fraction. The pellet was resuspended in a second solution (4 mM HEPES, 1 mM EDTA, pH = 7.4) and was centrifuged 2× at 12,000× *g* for 20 min. The new pellet was resuspended in a third solution (20 mM HEPES, 100 mM NaCl, 0.5% Triton X-100, pH = 7.2) for 1 h at 4°C and centrifuged at 12,000× *g* for 20 min. The supernatant collected corresponds to the non-PSD fraction (Triton-soluble). The remaining pellet was resuspended in a fourth solution (20 mM HEPES, 0.15 mM NaCl, 1% Triton X-100, 1% deoxycholicacid, 1% SDS, pH = 7.5) for 1 h at 4°C and was centrifuged at 10,000× *g* for 15 min to obtain a supernatant containing the PSD fraction (Triton-insoluble). The different fractions were then analyzed by WB.

### Quantification of synaptic connectivity

To quantify synaptic connectivity, we transducted primary hippocampal neurons in pre- and/or postsynaptic compartments of microfludic devices at DIV1 with lentiviruses carrying shNT and/or shFERMT2 (MOI = 10). At DIV14 cultures were fixed and immunostained against Synaptophysin I and Homer pre- and post-synaptic markers, respectively. Synaptic compartments were imaged with Zeiss LSM880 confocal microscope, using a 63× 1.4 NA objective and the AiryScan superresolution unit. Images were analyzed with Imaris software (Bitplane; Zürich, Switzerland) by reconstructing Synaptophysin I and Homer puncta in 3D. The volume and position information of all puncta were processed using a custom Matlab (MathWorks, Natick, MA) program. This program assigns each postsynaptic spot to the nearest presynaptic spot (within a distance threshold of 1 μm) and calculates the number of such assignments for all presynaptic puncta. The percentage of presynaptic spots not assigned by any postsynaptic spot was consistently used as a read-out of synaptic connectivity.

### Lentivirus injection

For stereotactic injections, C57Bl6/J mice (RRID:IMSR_JAX:000664) were anesthetized with 4% isoflurane (2 L/min) and placed in a stereotaxic frame (68528, RWD Life Science, Shenzhen, China) in which the head of the animal was fixed with a pair of ear bars and a perpendicular tooth bar. During surgical procedures 1.5% isoflurane (2 L/min) was delivered through a facial mask *via* spontaneous respiration. Their body temperature was maintained between 36.5 and 37.5°C with a homeothermic blanket. Head was shaved and Vetedine was applied. Wounds and pressure points were infiltrated with lidocaine. A skin incision was made along the sagittal midline of the scalp. Craniotomy was made to target the structures of interest. Lentiviruses were injected in right and left hippocampus (1.5 μL per hemisphere; 0.2 μL/min). After injections, wound clips were used for skin closure. For the sham group, surgical procedures were performed without any injection. During the surgery, the level of anesthesia was regularly verified by testing the nociceptive hind limb withdrawal reflex. Subjects were then allowed to recover in their home cages for at least 7 d before sacrifice for ex-vivo electrophysiological recordings.

### Hippocampal acute slices preparation

One week after the surgery, sagittal hippocampal brain slices were obtained using standard brain slicing methods. Mice were deeply anesthetized with isoflurane and decapitated. The brain was quickly removed and immersed in ice-cold pre-oxygenated artificial cerebrospinal fluid (aCSF) containing: 124 mM NaCl, 3.75 mM KCl, 2 mM MgSO_4_, 2 mM CaCl_2_, 26.5 mM NaHCO_3_, 1.25 mM NaH_2_PO_4_, and 10 mM Glucose and was continuously oxygenated (pH = 7.4; 27°C). 350 μm-thick slices were prepared using a Vibratome (VT 1000S; Leica Microsystems, Bannockburn, IL), and placed in a holding chamber filled with aCSF. Slices were allowed to recover in these conditions at least 1 h before recording.

### Electrophysiological recordings

For electrophysiological recordings, a single slice was placed in the recording chamber, submerged and continuously superfused with gassed (95% O_2_, 5% CO_2_) aCSF at a constant rate (2 mL/min). Extracellular fEPSPs were recorded in the CA1 stratum radiatum using a glass micropipette filled with aCSF. fEPSPs were evoked by the electrical stimulation of Schaffer collaterals/commissural pathway at 0.1 Hz with a glass stimulating electrode placed in the stratum radiatum (100 μsec duration).

To test the effect of miRNA-expressing lentiviruses on basal synaptic transmission, Input/Output (I/V) curves were constructed at the beginning of the experiment. The slope of fEPSPs was measured and plotted against different intensities of stimulation (from 0 to 100 μA).

Stable baseline fEPSPs were recorded by stimulating at 30% maximal field amplitude for 10 min prior to the beginning of the experiment (single-pulse stimulation every 10 s (0.1 Hz). The same intensity of stimulation was kept for the reminder of the experiment. For the paired-pulse facilitation (PPF) protocol, two stimulations were applied with 50, 100, 150, 200, 300, 400 and 500 ms interval. For long-term potentiation (LTP) protocol, after a 10 min stable baseline period, LTP was induced by the following stimulation protocol: 3 trains of 100 stimulations at 100 Hz at the same stimulus intensity, with 20 s intervals between trains. Following this conditioning stimulus, a 1 h test period was recorded where responses were again elicited by a single-pulse stimulation every 10 s (0.1 Hz) at the same stimulus intensity. Signals were amplified with an Axopatch 200B amplifier (Molecular Devices, Union City, CA) digitized by a Digidata 1550 interface (Axon Instruments, Molecular Devices, San Jose, CA) and sampled at 10 kHz. Recordings were acquired using Clampex (Molecular Devices) and analyzed with Clampfit (Molecular Devices). Experimenters were blinded to treatment for all experiments.

## RESULTS

### FERMT2 expression is dependent on miRNAs modulating APP metabolism

We used an unbiased screening approach to identify miRNAs that modulate APP metabolism in a HCS model that allows for the quantification of intracellular APP fragments (Chapuis et al. 2017). We screened a total of 2,555 mature human miRNAs in a 384-well plate format allowing us to identify 50 miRNAs (top and bottom 1%) with the strongest impact on APP metabolism (**Supplementary Table 2** and **Supplementary Fig. 2**). To determine which genes were potentially regulated by these top 50 hits, we selected the intersection of predictions resulting from at least four different algorithms (see methods) and thereby identified 6,009 putative miRNA-target genes. To further refine the list of predicted genes, we cross-checked them against a list of 832 genes that we previously identified to have a major impact on APP metabolism in a genome-wide siRNA screening (Chapuis et al. 2017) (**Supplementary Fig. 3**). This resulted in 180 common genes that are putative targets of 41 miRNAs. To determine if any of these 180 genes were preferentially regulated by this pool of 41 miRNAs, we performed 1 million drawing lots of 41 miRNAs among the 2,555 tested and compared them against the list of miRNAs predicted to bind in the 3’-UTR of each of the 180 genes (**Supplementary Fig. 3**). The AD genetic risk factor *FERMT2* (encoding Kindlin-2) was among the most significant genes (*p*-value < 2.77×10^−4^ after Bonferroni correction) that strongly modulate APP metabolism, and whose expression are potentially regulated by miRNAs that also strongly modulate APP metabolism. According to our screening, four miRNAs were predicted to target FERMT2 3’UTR: miR-582-5p, miR-200b-3p, miR-221-3p and miR-222-3p (**Fig. 1a**).

**Fig 2.**
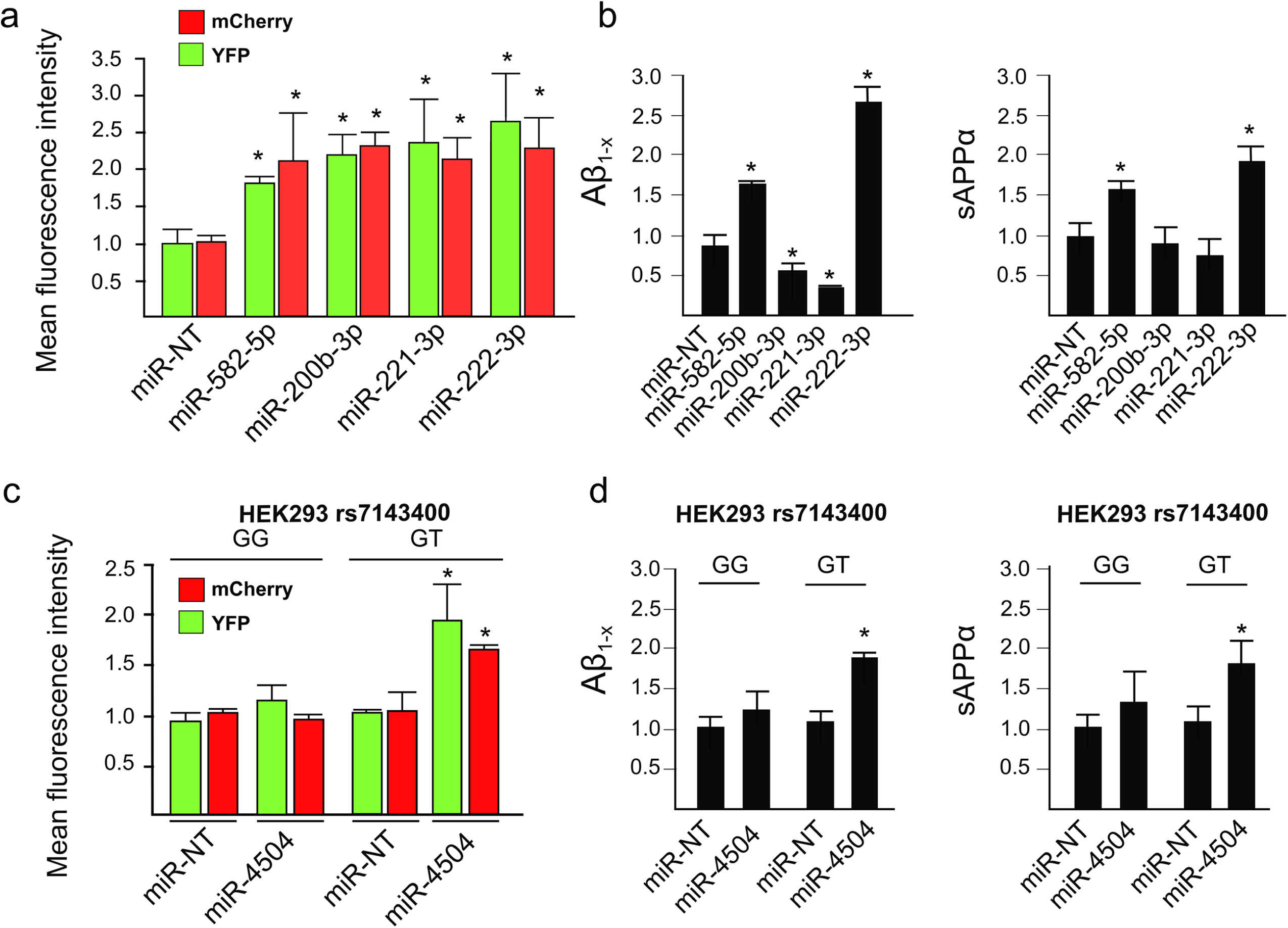
Validation of the effects of FERMT2-targeting miRNA on APP metabolism. **a**. Mean fluorescence intensity of intracellular mCherry and YFP signals obtained after miRNA transfection in HEK293 cells stably over-expressing a mCherry-APP^695WT^-YFP. **b**. Quantification of Aβ and sAPPα secretion after miRNAs transfection in HEK293 cells stably over-expressing a mCherry-APP^695WT^-YFP. **c**. Mean fluorescence intensity variation of intracellular mCherry and YFP signal obtained after miRNA transfection in HEK293^rs7143400-G/G^ or HEK293^rs7143400-G/T^ cell lines transiently over-expressing a mCherry-APP^695WT^-YFP. **d**. Quantification of Aβ and sAPPα secretion after miRNAs transfection in HEK293^rs7143400-G/G^ or HEK293^rs7143400-G/T^ cell lines transiently over-expressing a mCherry-APP^695WT^-YFP. Bar charts show mean ± SD. Mann–Whitney test; * *p* < 0.05.

**Fig. 3.**
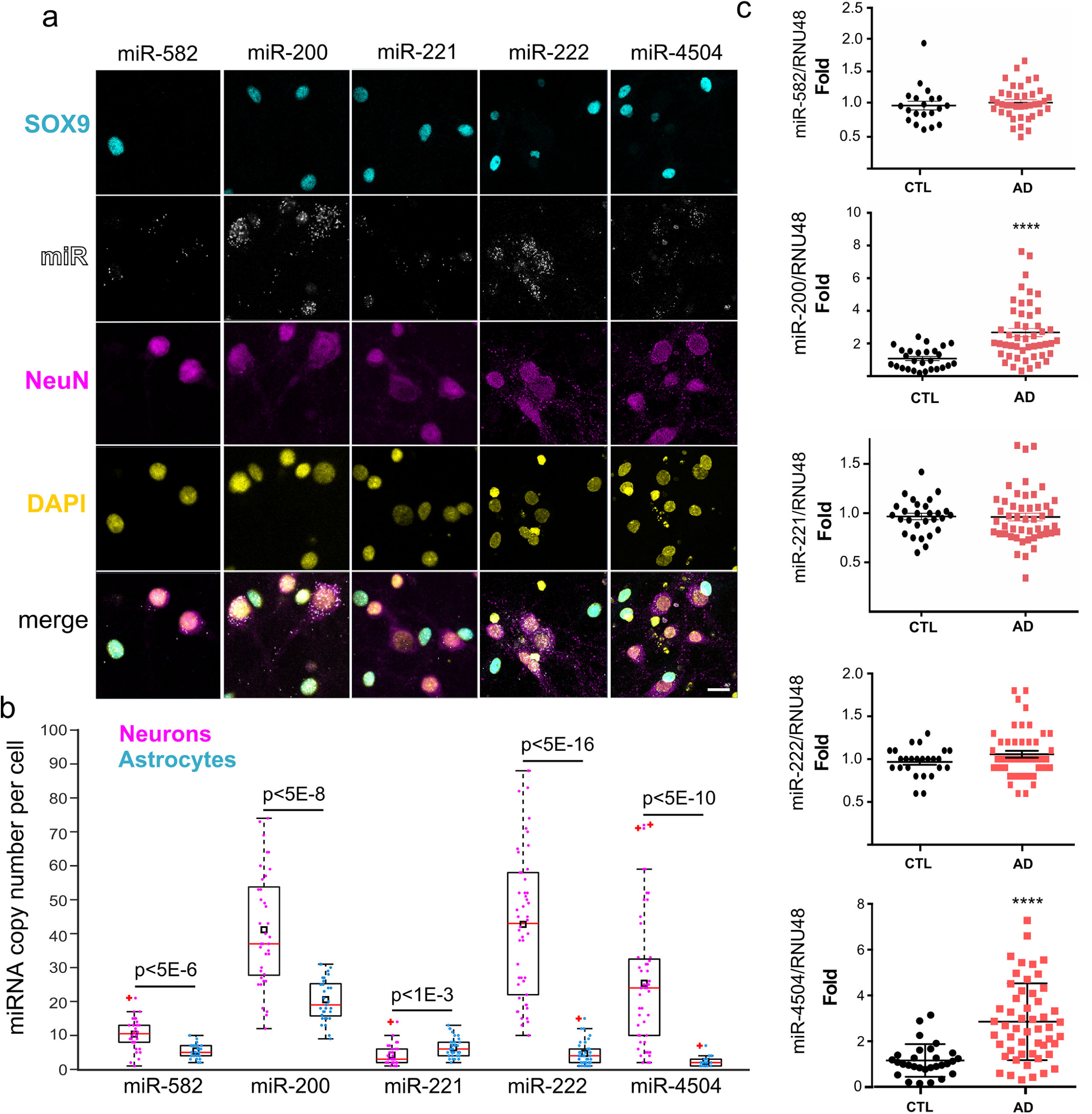
miRNA expression in primary neuronal cultures and in AD brains. **a**. Hybridization experiments in rat postnatal hippocampal neuronal cultures enabling single-copy detection of miRNA combined with immunocytochemistry against astrocytic (SOX9) and neuronal (NeuN) markers. Scale bar = 20 μm. The box plot shows the quantification of miRNA copy number in SOX9- or NeuN-positive cells (N > 30 cells for each condition). Black rectangles and red plus signs indicate sample mean and outliers, respectively. **b**. Relative miRNA expression levels in temporal lobes of non-demented (CTL) and AD groups. Mann–Whitney test; **** *p* < 0.0001.

### miRNA-dependent FERMT2 expression and genetic variation associated with AD risk

Our data indicate that regulation of the FERMT2 expression is dependent on miRNAs and we aimed to assess whether genetic variations associated with AD risk may modulate the miRNA-dependent expression of FERMT2. None of the variants localized within the FERMT2 3’-UTR were predicted to modify the binding of miR-582-5p, miR-200b-3p, miR-221-3p, or miR-222-3p to this region (Supplementary methods). In contrast, we had previously identified an AD-associated variant (rs7143400), where the minor T allele creates an 8-mer binding site for miR-4504 within the 3’-UTR of FERMT2 (Delay et al. 2016) (**Supplementary Fig. 4**). Supporting these predictions, we observed that miR-4504 led to reduced luciferase expression only in the presence of *FERMT2* 3’UTR rs7143400-T allele, whereas the four other miRNAs were able to induce a down-regulation regardless of the rs7143400 allele (**Fig 1b**).

**Fig 4.**
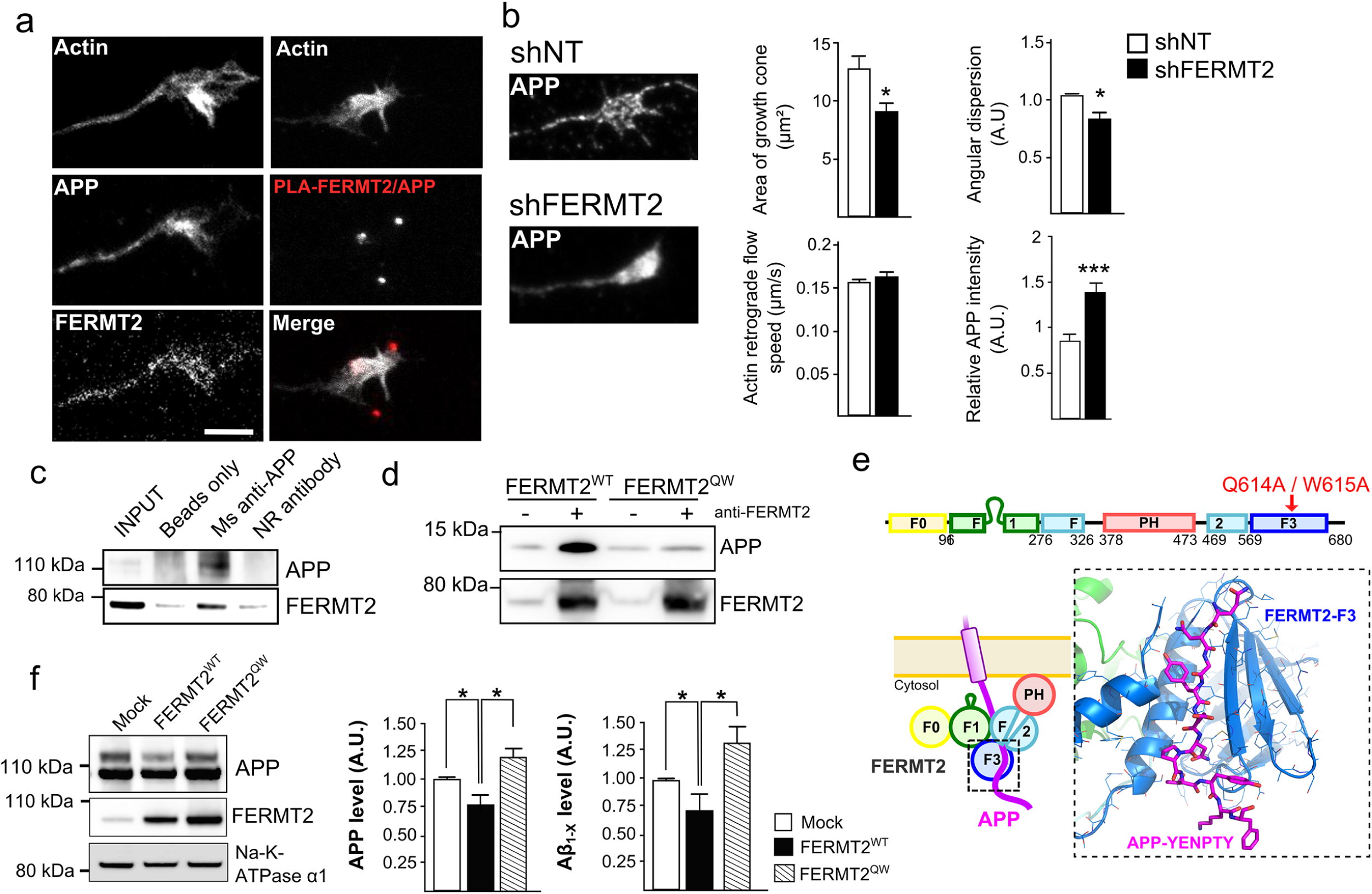
FERMT2 directly interacts with APP. **a**. Immunofluorescence images showing the presence of APP and FERMT2 within the axonal growth cone stained with SiR-Actin. The right panel shows the presence of PLA-FERMT2/APP puncta within the axonal growth cone. **b**. Impact of lentiviral transduction of non-targeting shRNA (shNT) or shRNA against FERMT2 (shFERMT2) on growth cone area, angular dispersion and speed of the actin retrograde flow, and APP immunostaining. **c**. Co-IP between endogenous APP and FERMT2 from membrane extracts of hippocampal PNC. Protein extracts were incubated with beads only, a mouse (Ms) antibody against APP (4G8) or a non-relevant (NR) antibody. **d**. APP pull-down experiment with wild type (WT) or mutated (QW) FERMT2. Protein extracts from HeLa cells overexpressing FERMT2^WT^ or FERMT^QW^ were incubated with recombinant APP C-terminal fragment (C100). **e**. The domain organization of FERMT2 protein (upper panel). Q614A/W615A (QW) mutation was reported to abolish the interaction of F3 domain of FERMT2 with the NxTY motif. The structural model of the FERMT2-APP complex (lower panel) was built by homology using the crystal structure of the FERMT2-Integrin-β3-tail complex (Li et al. 2017). **f**. The impact of FERMT2 on APP metabolism in HEK293-APP^695WT^ cells is reverted with the overexpression of FERMT2^QW^ compared to FERMT2^WT^. Scale bar = 5 μm. Mann–Whitney test; * *p* < 0.05.

We then assessed the impact of these five miRNAs on endogenous FERMT2 expression levels after their transfection in either HEK293^WT^ or rs7143400-mutated HEK293 cell lines (HEK293^rs7143400-G/T^) generated by CRISPR-Cas9 technology (**Fig. 1c** and **Supplementary Fig. 1 and 5**). Accordingly, transfection of miR-582-5p, miR-200b-3p, miR-221-3p, or miR-222-3p in HEK293 cells led to reduced FERMT2 expression whatever their genotype while transfection of miR-4504 decreased the endogenous FERMT2 expression only in the HEK293^rs7143400-G/T^ cell line (**Fig. 1d** and **e**). Similar effects were observed in HeLa cells that were genotyped to be heterozygous for rs7143400 (**Fig. 1c, d** and **e and Supplementary Fig. 5**).

**Fig 5.**
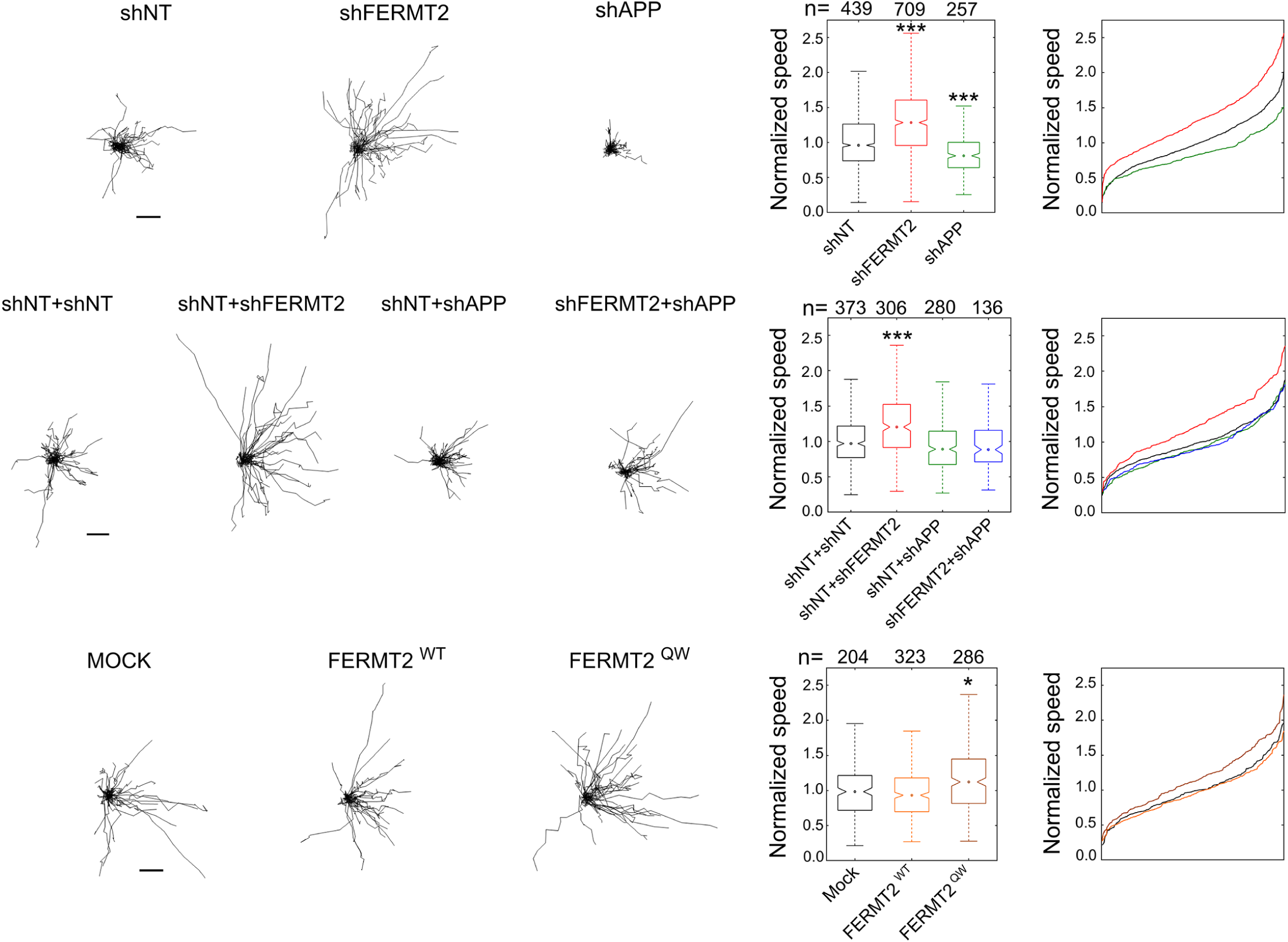
FERMT2 regulates axonal growth rate depending on APP expression. Impact of lentiviral transduction on axonal growth speed. Individual axon tracks from a representative set are plotted. Scale bars = 50 μm. Box plots and cumulative distribution plots are color-matched. n is the number of axons analyzed from at least three independent experiments. Kruskal-Wallis ANOVA with multiple comparisons; * *p* < 5×10^−3^; *** *p* < 5×10^−7^.

### Impact of miRNAs targeting FERMT2 on APP metabolism

These five miRNAs are thus potential candidates to modulate APP metabolism through a direct down-regulation of FERMT2. However, these miRNAs can also potentially target others genes strongly modulating APP metabolism (**Supplementary Table 3**). We reasoned that if a candidate miRNA affects APP metabolism mainly through down-regulating FERMT2, this candidate miRNA would have similar effects on APP metabolism as the direct FERMT2 down-regulation by siRNAs we had previously demonstrated, i.e. leading an increase of both intra- and extracellular byproducts of APP (Chapuis et al. 2017). To investigate this hypothesis, we used the data generated in our HCS approach (based on HEK293 cell line stably over-expressing a mCherry-APP695WT-YFP) in order to quantify intracellular byproducts of APP (Sannerud et al. 2011; Chapuis et al. 2017) and we also measured Aβ and sAPPα secretion after miR-582-5p, miR-200b-3p, miR-221-3p, or miR-222-3p transfections. Only miR-582-5p and miR-222-3p showed similar effects as FERMT2 down-regulation (Chapuis et al. 2017), *i*.*e*., they increased the levels of intracellular APP metabolites tagged by mCherrry and YFP and increased Aβ and sAPPα secretion (**Fig. 2a** and **b**).

Since the potential effects of miR-4504 would depend on the presence of the rs7143400 minor T allele, we were not able to test for its impact in our HCS model. We nevertheless took advantage of HEK293^rs7143400-G/T^ cells by co-transfecting them with miR-4504 and mCherry-APP^695WT^-YFP cDNA in order to mimic our HCS model. When compared to HEK293^rs7143400-G/G^, the transfection of miR-4504 in HEK293^rs7143400-G/T^ led to an accumulation of intracellular APP mCherry and YFP-tagged metabolites and an increase of Aβ and sAPPα secretion (**Fig. 2c** and **d**). MiR-4504 showed similar effects as FERMT2 down-regulation, and this observation further support that miR-4504 regulates APP metabolism as a function of the FERMT2 rs7143400 variant. In conclusion, we characterized that regulation of FERMT2 expression by miRNAs impacts APP metabolism, and potentially in a genetics-dependent manner.

### miRNA expression in different hippocampal cell types and in AD brains

To provide further physiological relevance to our findings, we first combined classical immunocytochemistry with RNA hybridization which allows for the detection of miRNAs at single-copy sensitivity. We observed that miR-200, miR-222 and miR-4504 were mainly expressed in neurons when compared to astrocytes (**Fig. 3a**). We next measured the expression levels of these miRNAs in the post-mortem brain samples from 52 AD patients and 30 control subjects. We observed that the expression levels of miR-200 and miR-4504 were significantly higher in AD brains than in controls (**Fig. 3b**). Collectively, these data suggest that endogenous FERMT2 expression and its impact on APP metabolism are dependent on the expression of several miRNAs, two of which are over-expressed in the brains of AD cases and, among these two, one impacts APP metabolism in the presence of a genetic variant associated with AD risk.

### Pathway analyses suggest FERMT2/APP interaction to be involved in axonal growth

Little is known about the physiological processes that require the regulation of APP expression and/or its metabolism by miRNAs. To obtain a list of potential physiological pathways to be further investigated, pathway enrichment analysis was performed using the 41 candidate miRNAs that strongly modified APP metabolism in our HCS (**Supplementary Table 2**). This analysis revealed that the candidate miRNAs are predicted to regulate neuronal pathways, such as axonal guidance (**Table 1a**). Since these 41 miRNAS potentially target 180 genes that strongly modulate APP metabolism (**Supplementary Table 3**), we also performed pathway-enrichment analysis using these 180 genes. This analysis revealed that these genes are predicted to be involved in axonal guidance among others (**Table 1b**). In conclusion, both miRNAs and genes modulating APP metabolism, *e*.*g*., FERMT2, potentially play a role in axonal guidance.

APP is already known to be enriched in axonal growth cones during nervous system development and acts as a co-receptor for axon guidance and cell migration cues through its interaction with the extracellular matrix (Soldano & Hassan 2014) (Sosa et al. 2013). We thus investigated the potential involvement of FERMT2 in axonal growth. Using primary neurons cultured in microfluidic devices that fluidically isolate axons from their cell bodies, we first observed the co-localization of endogenous FERMT2 with APP in the growth cones (**Fig. 4a**). We then addressed the impact of FERMT2 silencing on axonal growth cone morphology using lentiviral vectors expressing either shRNA against FERMT2 (shFERMT2) or a non-targeting shRNA (shNT). Actin staining revealed that FERMT2 under-expression led to a significant decrease in growth cone area (9.13±0.71 vs 12.79±1.10 μm^2^), as well as in the angular dispersion of growth cone filopodia during axonal growth (0.67±0.04 vs 0.84±0.02) (**Fig. 4b and Supplementary Fig. 6**). Of note, no significant impact on actin retrograde flow rate was observed (0.166±0.003 vs 0.157±0.002 μm/s). These observations suggest a potential impairment of the exploration behavior of the growth cones due to FERMT2 silencing, but not an effect on actin dynamics *per se*. FERMT2 under-expression was also associated with an accumulation of endogenous APP in the growth cones (1.38±0.11 vs 0.85±0.08, after normalization by the growth cone area).

**Fig 6.**
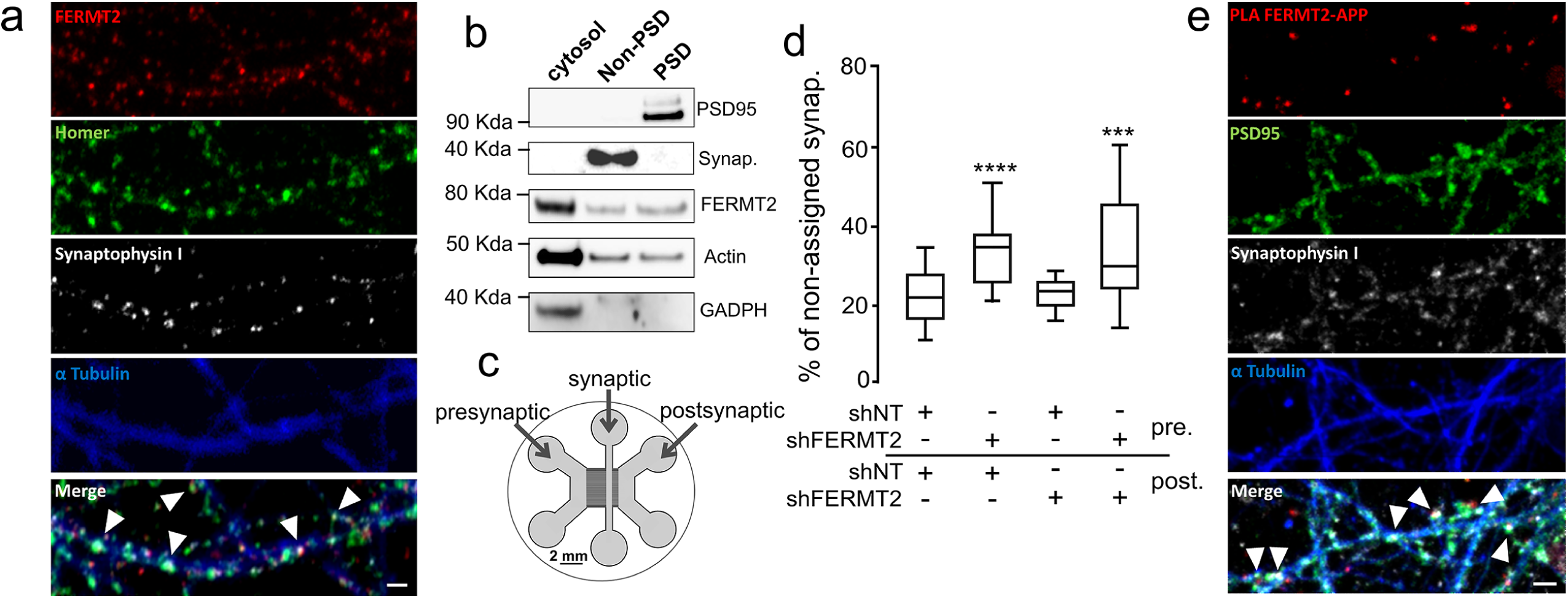
FERMT2 is present at the synapse and controls synaptic connectivity. **a**. Immunofluorescence in hippocampal primary neuronal culture showing the co-localization of FERMT2 puncta with pre- and postsynaptic markers, Synaptophysin and Homer, respectively. **b**. Synaptic fractionation experiment revealed the presence of FERMT2 in both pre- and postsynaptic compartments. **c**. Schematics of the tricompartmental microfluidic device. The use of microchannels with different lengths ensures that only axons arrive from the presynaptic to the synaptic compartment, where synapses can be observed independently of the cell bodies. The device also permits lentiviral transductions to be performed exclusively in the pre- and postsynaptic compartments. **d**. Synaptic connectivity as a function of FERMT2 under-expression in pre- and postsynaptic chambers. Increased fraction of Synaptophysin spots not assigned by a Homer spot within a distance threshold of 1 μm is indicative of decreased synaptic connectivity. **e**. PLA-FERMT2/APP puncta were observed at the synapses stained for pre- and postsynaptic markers. Scale bars = 2 μm.

By performing proximity ligation assay (PLA), we observed PLA-FERMT2/APP signals in axonal growth cones (**Fig. 4a**), suggesting a potential function of the FERMT2/APP complex in axon growth behavior. To address this, we first tested the possibility that APP and FERMT2 form a protein-protein complex *via* three complementary approaches: (i) Pull-down of endogenous APP from hippocampal primary neuronal culture extracts co-immunoprecipitated endogenous FERMT2 (**Fig. 4c**). (ii) Over-expression of FERMT2^WT^ was also able to pull-down the recombinant intracellular domain of APP (**Fig. 4d**). (iii) In addition, we generated a Q_621_W_622_AA FERMT2 mutant (FERMT2^QW^) which was previously shown to abolish the interaction between the FERMT2 F3 domain and the NxTY motif of Integrin-β3 (which is also present within the intracellular domain of APP) (Ma et al. 2008). Remarkably, when over-expressed in HEK293 cells, FERMT2^QW^ was not able to pull-down the recombinant intracellular domain of APP. Cumulatively, these findings support a direct interaction between FERMT2 and APP. Based on the recently solved crystal structure of FERMT2 in complex with the integrin-β3-tail (Li et al. 2017), we built a structural model of the FERMT2/APP complex (**Fig. 4e**), supporting our hypothesis that a protein-protein interaction exists between FERMT2 and APP.

We next assessed the biological impact of the FERMT2/APP interaction on APP metabolism. By performing extracellular biotinylation experiments, we observed that FERMT2 over-expression in HEK293-APP^695wt^ cell line decreased the levels of APP at the cell surface, an effect that was abolished by the presence of the QW mutation (**Fig. 4f**). Further, a dominant negative effect of the FERMT2^QW^ mutant was observed: its over-expression impacted APP metabolism similarly to FERMT2 silencing, *i*.*e*., resulting in increased mature APP at the cell surface and increased Aβ production, as previously reported (Chapuis et al. 2017). Altogether, our data suggest that a FERMT2/APP interaction is necessary for FERMT2 to have an impact on APP metabolism.

In order to characterize in-depth the impact of FERMT2 and/or APP expression on axonal growth, we conducted time-lapse microscopy and measured axon growth speed at DIV5 following lentiviral transduction (shNT, shFERMT2, or shAPP) of neurons in microfluidic devices at DIV1. FERMT2 silencing led to 31.7% increase in axon growth speed (**Fig. 5 and SupplementaryFig. 6**). Conversely, APP under-expression led to 16.7% decrease in axon growth speed. Remarkably, silencing of APP was able to fully abolish the effect of FERMT2 under-expression on axon growth speed, suggesting that APP was required for the molecular mechanism by which FERMT2 controls the axon growth speed. In addition, we observed that FERMT2^QW^ mutant over-expression was able to induce 15.9% increase in axon growth speed (**Fig. 5**). Since over-expression of FERMT2^WT^ did not show any impact, these data also suggested a potential dominant negative effect of the FERMT2^QW^ mutant and further supported the involvement of FERMT2/APP complex in axonal growth.

### FERMT2 is present at the synapse and controls synaptic connectivity

Next, we investigated the impact of FERMT2 silencing on neuronal maturation at DIV14. First, co-staining between FERMT2 and synaptic markers (Synaptophysin and Homer) suggested the localization of FERMT2 at the synapse (**Fig. 6a**). The presence of FERMT2 in both pre- and postsynaptic compartments was confirmed by synaptosomal purification (**Fig. 6b**). To control shRNA expression separately in pre- or postsynaptic neurons, hippocampal neurons were cultured in microfluidic devices that promote synapse formation in an isolated chamber (Taylor et al. 2010). Thanks to the use of narrow microchannels, these devices spatially isolate neurites from their cell bodies and allow lentiviral transductions to be conducted in different compartments, thereby allowing us to silence FERMT2 expression at the pre- and/or postsynaptic levels (**Fig. 6c and Supplementary Fig. 6**). The effects of shRNA expression (DIV1) on synaptic connectivity were assessed by confocal microscopy of synaptic markers (DIV14) followed by three-dimensional image segmentation and quantification. Under-expression of FERMT2 in the pre-synaptic chamber led to a decrease in synaptic connectivity, whereas no such effect was observed when under-expressing FERMT2 in the post-synaptic compartment (**Fig. 6d**). Altogether, our data suggest that FERMT2 expression is required for synapse connectivity. Moreover, PLA-FERMT2/APP signals were co-localized with Synaptophysin and Homer puncta (**Fig. 6e**), supporting the possibility of the involvement of the FERMT2/APP complex in synapses.

### FERMT2 expression regulates synaptic plasticity in an APP dependent manner

We sought to establish the functional impact of FERMT2 and/or APP silencing on paired-pulse facilitation (PPF) and long-term potentiation (LTP) in *ex vivo* mouse (10-week-old male) hippocampal slices, after stereotactic lentivirus injection allowing for the expression of shNT, shFERMT2, shAPP, or shFERMT2+shAPP.

Broadly speaking, PPF arises due to increased presynaptic Ca^2+^, which leads to the release of neurotransmitter in two distinct waves. In this situation, two action potentials in the presynaptic cell produce two excitatory postsynaptic potentials (EPSPs) in the postsynaptic cell: the first action potential produces a first EPSP, but the second action potential produces an EPSP that is larger than the EPSP produced by the first. PPF modulation therefore highlights a modulation in presynaptic neurotransmitter release (Gebhardt et al. 2019). Using this read-out as a proxy for presynaptic function, we observed a significant decrease in PPF in shFERMT2-infected mice compared to shNT-infected control mice (**Fig. 7a**). This PPF impairment however was rescued when APP was also down-regulated (shAPP+shFERMT2 group).

**Fig 7.**
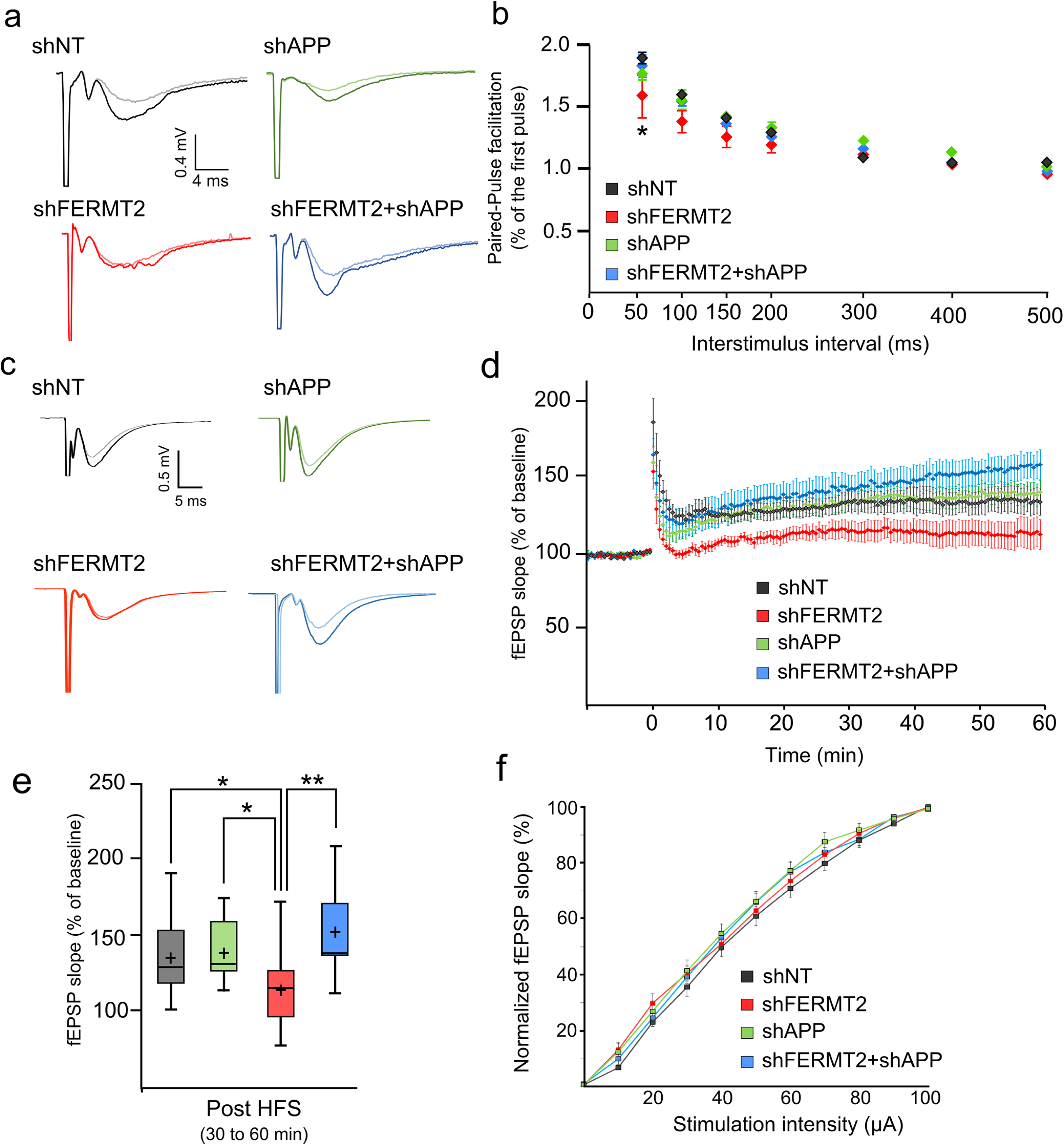
FERMT2 under-expression alters PPF and LTP in an APP-dependent manner. **a**. Paired-pulse facilitation (PPF) as a function of the interstimulus interval 7 days after viral injection of indicated lentivirus. N = 3 mice; 2 slices per animal. **b**. Exemplary fEPSP traces during baseline (light line) and 30-60 min after LTP induction (dark line). **c**. Time course of the average slope of elicited fEPSP responses following LTP induction by a tetanic stimulation protocol in hippocampal CA1 synapses after viral injection. Time-point 0 represents the delivery of the tetanic stimulation. Slopes of each fEPSP are normalized by the baseline and plotted against time. **d**. Box plots of the average slope response during 30-60 min post LTP induction. HFS: High frequency stimulation. N = 5 mice; 2 slices per animal. **e**. Normalized average slope of fEPSP evoked in hippocampal slices from animals injected with the indicated lentivirus. Recordings have been performed in the stratum radiatum of hippocampal CA1 region with electrical stimulation of Schaffer collaterals (see Methods). Unpaired t-test; * *p* < 0.05; ** *p* < 0.01.

In separate experiments, tetanic stimulation was delivered to the Shaffer collaterals (SC) in order to induce LTP in hippocampal slices (**Fig. 7b**). Tetanic stimulation of the SC resulted in a robust, long-lasting potentiation of the field excitatory postsynaptic potential (fEPSP) slope in slices from mice infected with shNT and with shAPP, whereas LTP was impaired in slices from shFERMT2-infected mice. This suggests that an LTP deficit was observed in hippocampal slices infected with shFERMT2, but not in those infected with shAPP (**Fig. 7c** and **7d**). Remarkably, this deficit was abolished when both APP and FERMT2 were silenced, suggesting that APP was required for the molecular mechanism by which FERMT2 impacts LTP.

Importantly, in these slices, no significant difference was observed for the normalized average slope of the evoked fEPSP, indicating no alteration of the CA1 basal synaptic transmission occurred in any of the groups analyzed (**Fig. 7e, Supplementary Fig. 7**).

Altogether, these data are in agreement with our previous observations that FERMT2 is involved in the pre-synaptic compartment and modulates synaptic connectivity in an APP-dependent manner.

## DISCUSSION

As in other multifactorial diseases, GWAS in AD are agnostic approaches, and how a genetic risk factor is implicated in pathophysiological processes is typically unknown. Sometimes, even the physiological functions of an AD genetic risk factor in the brain are not known. Understanding the role of these genes is thus a challenge that requires several key questions to be addressed: (i) Does the corresponding protein interact (directly or indirectly) with other key players and pathways known to be involved in AD? (ii) What is (are) the functional variant(s) responsible for the GWAS signal and does this (do these) variant(s) impact the biological function of the corresponding protein and its interaction with key players of AD?

To answer these questions, we developed systematic approaches to determine the genes that are involved in APP metabolism, a major player in AD development. To this end, we had previously developed an HCS, based on the quantification of intracellular APP fragments, to measure the impact of under-expression of 18,107 genes (*via* siRNA pools) on APP metabolism (Chapuis et al. 2017). In the current study, we screened the impact of the over-expression of 2,555 miRNAs on APP metabolism with the hypothesis that genes (i) that modulate the APP metabolism and (ii) whose expression levels are regulated by miRNAs that also modulate the APP metabolism are likely some of the key actors controlling the APP metabolism and functions. The convergence of these two agnostic screens highlighted FERMT2, a GWAS-defined genetic risk factor of AD, for which almost nothing is known in the cerebral and AD contexts.

We demonstrated that a direct interaction between FERMT2 and APP –through the F3 domain of FERMT2 and the NxTY motif within APP’s intracellular domain– is necessary for FERMT2 to have an impact on APP metabolism. Moreover, we observed that the FERMT2/APP interaction could be involved in the regulation of axonal growth, in line with APP’s function within the growth cone (Sosa et al. 2013) (data we replicated in this study). It has been reported that FERMT2 is required for the recruitment and activation of focal adhesion kinase and the triggering of integrin signaling (Theodosiou et al. 2016). In neurons, the focal adhesion pathway is involved in synaptic density and activity through regulating the dendritic spine shape, stability, and the signalling machinery therein (Hotulainen & Hoogenraad 2010). That is why we also analyzed synaptic plasticity, a read-out highly relevant to AD, where synaptic dysfunction/loss is one of the earliest events observed. FERMT2 under-expression had detrimental effects on PPF (presynaptic) and LTP (postsynaptic). Remarkably, in both cases, the detrimental effect of FERMT2 under-expression was dependent on APP expression. In this context, it is important to note that numerous evidence indicate that presynaptic physiological functions involving APP, which has been recently proposed as a structural and functional regulator of the hippocampal presynaptic active zone (Weingarten et al. 2017), could be major molecular players in AD (Barthet & Mulle 2020). As FERMT2 silencing leads to an accumulation of full-length APP and all its by-products (including Aβ peptides), we can hypothesize that these accumulations could be involved in the synaptic dysfunction observed due to FERMT2 under-expression, although further experiments are needed to decipher the potential causal link between FERMT2 and APP, *i*.*e*., to determine whether FERMT2 impacts the function of full-length APP or invokes Aβ synaptotoxicity. This is of particular interest, since APP shedding strongly enhances its cell adhesion and synaptogenic activity (Stahl et al. 2014). Moreover, APP’s intracellular domain is required for normal synaptic morphology and synaptic plasticity, suggesting that its intracellular interaction partners could be required for proper synaptic function (Klevanski et al. 2015). Remarkably, we have recently proposed a circular model of AD pathogenesis, where the core of the focal adhesion pathway –which FERMT2 and APP are part of– may participate in the dysfunction of synaptic plasticity in AD (Dourlen et al. 2019).

We have also identified that *FERMT2* expression level is highly regulated by miRNAs which could be preferentially expressed in neurons. In addition, we previously identified the rs7143400 variant located in *FERMT2* 3’UTR to be susceptible to alter a binding site for miR-4504 (Delay et al. 2016). Here, in addition to *in silico* prediction suggesting the impact of this variant on miRNA binding (**Supplementary Fig. 4**), we demonstrate that this variant is functional: the AD-associated rs7143400 T allele down-regulates *FERMT2* and modulates APP metabolism *via* its interaction with miR-4504. Remarkably, we observed that miR-4504 is over-expressed in the brains of AD cases compared to controls, and is mainly expressed in neurons in primary mixed hippocampal cultures.

Together, our data indicate that a deleterious over-expression of miR-4504 can lead to a decrease in FERMT2 expression in individuals bearing the rs7143400 minor T allele, which subsequently modulates APP metabolism. Interestingly, similar mechanism has been reported for genetic variant associated with AD risk in APP 3’UTR which regulates APP expression through miRNA binding (G et al. 2016). Supporting a link between FERMT2 and APP metabolism, studies from cohorts of patients have reported an association between variants in *FERMT2* gene and Aβ in CSF (Chapuis et al. 2017) and brain amyloidosis (Apostolova et al. 2018).

Here, we propose that FERMT2 down-regulation at the earliest stage of AD would depend in part on (i) the miR-4504 expression, (ii) cerebral cell type (*i*.*e*., neurons), and (iii) the presence of the rs7143400 minor T allele (observed in 9% of Caucasians). Unfortunately, it is important to keep in mind that all these constraints will make difficult, if not impossible, to detect such a miRNA-dependent decrease in FERMT2 mRNA levels. Of note, this point may also underline the limitation of expression databases in deciphering the mechanisms underlying the functional effects of GWAS variants, for they do not allow capturing (even hide) subtle mechanisms.

In publicly-available RNA-seq analyses (Mayo Clinic Brain Bank), an over-expression of FERMT2 mRNA has been observed in post-mortem human temporal cortex of AD patients relative to healthy controls (Sullivan et al. 2018). Even though a small sample size did not allow us to observe such a variation in FERMT2 mRNA levels, we nevertheless detected an increase in FERMT2 protein levels as a function of Braak stage, especially at later stages (**Supplementary Fig. 8)**. This point is of particular importance since in the Genotype-Tissue Expression Database (GTEx Consortium et al. 2015), FERMT2 variants associated with an increase in AD risk at the genome-wide significance level are also part of an expression quantitative trait locus, significantly associated with decreased brain expression of FERMT2 mRNA (sentinel variant in GWAS rs17125924; −18%; *p*-value = 2×10^−6^). Of note, there is a strong linkage disequilibrium between rs7143400 and the GWAS hit rs17125924 (R^2^ = 0.78) and rs7143400 has a lower Minor Allele Frequency (MAF = 0.09) and subsequently lower association (*p* = 7.14×10^−5^) than rs17125924 (MAF = 0.20; *p* = 6.6×10^−7^). Altogether, these results strongly support the notion that FERMT2 down-regulation is deleterious at the earliest stages of the disease, whereas FERMT2 over-expression may occur as a pathological consequence at a later stage. In conclusion, we propose that FERMT2 under-expression through miRNAs and/or genetic regulation leads to synaptic dysfunction in an APP-dependent manner. Our hypothesis may thus call for new therapeutic approaches in AD targeting FERMT2 and/or APP function, rather than Aβ peptide production/clearance.

## Acknowledgements

F. E. benefited from a doctoral grant co-funded by Hauts-de-France Regional Council and Institut Pasteur de Lille. The authors thank the BICeL and EquipEx ImagInEx HCS platforms of the Institut Pasteur de Lille and Alexandre Vandeputte for technical assistance. The authors thank Karine Blary at the IEMN Lille for the microfabrication work. This work was partly supported by the French RENATECH network. This study was funded by INSERM, Institut Pasteur de Lille, the Centres of Excellence in Neurodegeneration-ANR program (CoEN, GWAS in AD: focus on microRNA), the Canadian Institute of Health Research (CHIR), la Fondation Alzheimer (Syn-Alz), the EU Joint Programme – Neurodegenerative Diseases Research (JPND; 3DMiniBrain) and Fondation Vaincre Alzheimer (FR-17006p). This work was also funded by the Lille Métropole Communauté Urbaine and the French government’s LABEX DISTALZ program (development of innovative strategies for a transdisciplinary approach to Alzheimer’s disease). This work was supported for M.H by Academy of Finland (grant number 307866), Sigrid Jusélius Foundation and the Strategic Neuroscience Funding of the University of Eastern Finland. The authors thank the vectorology platform Transbiomed for lentivirus production.

## Author contributions

J.-C. L. and J. C. designed and supervised research. A. F., F. E., F. C. and C. B. performed APP metabolism and FERMT2/APP interaction analyses. X. H. developed in silico model for FERMT2/APP interaction. F. E. and D. K. performed and analyzed axon growth experiments. A. C., A. F., J. D. and J. C. developed Crispr/Cas9 model and/or performed subsequent analyses. C. D., A.-C. V., A. F. and B. G.-B. designed and/or performed miRNA screening and/or statistical analyses. S. H. and M. F. performed and analyzed electrophysiology experiments. A. F., T. M., F. D. and S. D. performed primary neuronal cultures. M. M., M. T., I. P. and M. H. analyzed transcriptomic and/or proteomic data of FERMT2 expression in brains. E. B. and S. S. H. performed miR expression quantification in brains. F. E., N. M., D. K. and J. C. participated in image acquisition and analyses of APP/FERMT2 interaction and/or synapse density. F. E., P. A., J. D., D. K., J.-C. L. and J. C. wrote and/or revised the paper.

