## supplementary data for "Alzheimer’s genetic risk factor *FERMT2* (Kindlin-2) controls axonal growth and synaptic plasticity in an APP-dependent manner"

### Supplementary Methods

#### High-content miRNA screening

The miRNA library (2,555 miRNA mimics, mirVana™ library, Ambion, Austin, TX) was screened in HEK293 cells stably over-expressing mCherry-APP695$^{wt}$-YFP (**Supplementary Fig. 2**) according to a previous protocol (Chapuis et al. 2017). Briefly, each miRNA (50 nM) was first transferred to 384-well with D-PBS containing 0.1 µl of Lipofectamine RNAiMax (Life Technologies). Then approximately 3,000 HEK293-mCherry-APP695wt-YFP cells were distributed in each well and incubated for 3 d at 37°C. After removal of the cell medium and Hoechst staining, cells were fixed using 10% formalin before image acquisition using an InCell Analyser 6000 (GE Healthcare, Chicago, IL) high-resolution automated confocal microscope.

A customized image analysis software (Columbus 2.7; PerkinElmer, Villebon-sur-Yvette, France) was used for the image analysis and quantification as reported previously (Chapuis et al. 2017). The mean fluorescence intensity of each signal was normalized with the fold change induced by the non-targeting miRNA in the same plate. To evaluate the impact of each miRNA, an average of 1,000 cells were analyzed per run (n = 3). For quality control, we used strictly standardized mean difference with β-score > 3 based on two separate controls (siRNA-APP and miR-NT, **Supplementary Fig. 2**).

#### *In silico* prediction of miRNA targets and functions

Seven publicly-available algorithms were used for the prediction of miRNA-targeted genes in the brain: DIANA-microT-CDS (v5.0, r21) (Paraskevopoulou et al. 2013), MiRanda (August 2010 release) (John et al. 2004), mirDB (v5.0) (Wong & Wang 2015), miRTarBase (v6.1) (Hsu et al. 2011), rna22 (v2.0) (Miranda et al. 2006), TargetMiner (May 2012 release) (Bandyopadhyay & Mitra 2009), and TargetScan (v7.1) (Agarwal et al. 2015). In order to minimize false-positive predictions, miRNA-targeted genes were defined as genes predicted by at least 4 out of the 7 algorithms used (**Supplementary Table 2**). To predict the function(s) of miRNAs that target these genes, pathway enrichment analysis was performed using DIANA Tools mirPath (v3.0) (Vlachos et al. 2015). Pathway analysis of targeted genes was performed using KEGG PATHWAY Database.

#### Human brain samples for FERMT2 expression (mRNA and protein)

The neuropathological cohort consists of *post-mortem* temporal cortical brain samples from 71 individuals. A detailed description of cohort validation and sample and data processing are described in (Marttinen et al. 2019). Briefly, at autopsy, assessment of the extent of AD-related neurofibrillary pathology was performed by immunostaining of paraffin sections with the AT8 antibody (Thermo Fisher Scientific, RRID:AB_223647), detecting hyperphosphorylated tau (Braak et al. 2006). The cohort was divided into seven disease severity stages based on Braak staging: Braak stages 0-VI. 60 of the 71 samples underwent microarray analysis and 36 proteomic analysis, with a total of 25 samples undergoing both microarray and proteomic analysis. The study was approved by the Ethics Committee of Kuopio University Hospital, University of Eastern Finland, the Finnish National Supervisory Authority, and the Finnish Ministry of Social Affairs and Health (License number 362/06.01.03.01/2017)

**Supplementary Table 1.**

Characteristics of human cohort used for miRNA quantification in human brain samples

| **Characteristics** | **Ctrl** | **AD** |
|---|---|---|
| n | 30 | 52 |
| Male % | 67.74 | 50.98 |
| Mean age of death | 74.86 | 78.34 |
| Post-mortem delay (hours) | 16.28 | 17.65 |
| Braak score I/II/III/IV/V/VI | 4/3/0/0/0/0 | 0/0/10/13/14/15 |

**Supplementary Table 2.** Variation of YFP and mCherry signals (Log2 of fold change) for the best miRNA hits exhibiting the strongest variations. Aβ secretion was also quantified in medium. SD: standard deviation. ND: Not detected.

| | | YFP (Log2) | | mCherry (Log2) | | $A\beta_{1-x}$ | |
|---|---|---|---|---|---|---|---|
| **miRNA** | **Refseq** | Mean Fold_change | SD | Mean Fold_change | SD | Mean Fold_change | SD |
| hsa-miR-1343-3p | MIMAT0019776 | -0,788 | 0,051 | -1,061 | 0,133 | ND | - |
| hsa-miR-16-1-3p | MIMAT0004489 | 1,018 | 0,067 | 1,242 | 0,051 | 5,218 | 1,596 |
| hsa-miR-181b-3p | MIMAT0022692 | -0,727 | 0,254 | -1,250 | 0,402 | 0,940 | 0,236 |
| hsa-miR-190a-3p | MIMAT0026482 | -0,800 | 0,150 | -1,075 | 0,172 | 1,261 | 0,046 |
| hsa-miR-193a-3p | MIMAT0000459 | 1,353 | 0,209 | 1,153 | 0,054 | 13,186 | 0,159 |
| hsa-miR-193b-3p | MIMAT0002819 | 1,662 | 0,104 | 1,292 | 0,129 | 3,975 | 0,113 |
| hsa-miR-194-3p | MIMAT0004671 | -1,091 | 0,239 | -1,595 | 0,037 | 1,263 | 0,268 |
| hsa-miR-200b-3p | MIMAT0000318 | 1,151 | 0,181 | 1,205 | 0,103 | 0,901 | 0,151 |
| hsa-miR-2115-5p | MIMAT0011158 | 1,574 | 0,151 | 1,270 | 0,326 | ND | - |
| hsa-miR-221-3p | MIMAT0000278 | 1,238 | 0,328 | 1,083 | 0,192 | 0,559 | 0,007 |
| hsa-miR-222-3p | MIMAT0000279 | 1,391 | 0,361 | 1,180 | 0,260 | 4,392 | 0,286 |
| hsa-miR-3160-3p | MIMAT0015034 | 1,675 | 0,285 | 1,457 | 0,228 | 2,549 | 0,434 |
| hsa-miR-3160-5p | MIMAT0019212 | 1,465 | 0,169 | 1,509 | 0,163 | ND | - |
| hsa-miR-3179 | MIMAT0015056 | 0,805 | 0,386 | 1,101 | 0,069 | 0,761 | 0,078 |
| hsa-miR-3188 | MIMAT0015070 | -0,779 | 0,064 | -1,066 | 0,083 | 2,385 | 0,427 |
| hsa-miR-32-3p | MIMAT0004505 | -0,864 | 0,203 | -1,249 | 0,082 | 0,496 | 0,042 |
| hsa-miR-3529-5p | MIMAT0019828 | -1,064 | 0,154 | -0,932 | 0,050 | -0,048 | 0,105 |
| hsa-miR-3616-5p | MIMAT0017995 | 2,284 | 0,400 | 1,463 | 0,016 | ND | - |
| hsa-miR-3663-3p | MIMAT0018085 | 0,969 | 0,470 | 1,166 | 0,153 | ND | - |
| hsa-miR-3668 | MIMAT0018091 | -0,567 | 0,260 | -0,976 | 0,082 | ND | - |
| hsa-miR-3977 | MIMAT0019362 | 1,285 | 0,793 | 0,967 | 0,668 | ND | - |
| hsa-miR-4429 | MIMAT0018944 | 1,172 | 0,233 | 1,050 | 0,193 | ND | - |
| hsa-miR-4477a | MIMAT0019004 | -0,354 | 0,172 | -1,975 | 1,370 | ND | - |
| hsa-miR-4773 | MIMAT0019928 | 1,819 | 0,274 | 1,540 | 0,163 | 11,263 | 0,477 |
| hsa-miR-488-5p | MIMAT0002804 | -0,927 | 0,243 | -1,137 | 0,071 | 0,120 | 0,077 |
| hsa-miR-506-3p | MIMAT0002878 | -0,988 | 0,310 | -1,015 | 0,124 | 0,584 | 0,053 |
| hsa-miR-511-3p | MIMAT0026606 | -0,778 | 0,164 | -1,077 | 0,204 | ND | - |
| hsa-miR-513c-3p | MIMAT0022728 | -0,833 | 0,041 | -1,097 | 0,081 | ND | - |
| hsa-miR-5193 | MIMAT0021124 | -0,991 | 0,042 | -1,145 | 0,116 | 0,012 | -0,110 |
| hsa-miR-5579-5p | MIMAT0022269 | -0,983 | 0,119 | -0,953 | 0,325 | ND | - |
| hsa-miR-5582-5p | MIMAT0022279 | -0,709 | 0,061 | -1,047 | 0,135 | 0,267 | 0,025 |
| hsa-miR-582-5p | MIMAT0003247 | 0,865 | 0,058 | 1,043 | 0,443 | 2,714 | 0,004 |
| hsa-miR-6783-3p | MIMAT0027467 | -0,866 | 0,149 | -1,141 | 0,085 | 0,414 | 0,290 |
| hsa-miR-6884-5p | MIMAT0027668 | -0,769 | 0,207 | -1,042 | 0,178 | 0,592 | 0,088 |
| hsa-miR-7151-3p | MIMAT0028213 | 1,772 | 0,197 | 1,545 | 0,070 | 5,990 | 0,109 |
| hsa-miR-7160-3p | MIMAT0028231 | 0,904 | 0,116 | 1,208 | 0,084 | 5,140 | 0,542 |
| hsa-miR-7160-5p | MIMAT0028230 | 1,120 | 0,257 | 1,134 | 0,167 | 0,414 | 0,290 |
| hsa-miR-7-5p | MIMAT0000252 | -0,819 | 0,014 | -1,027 | 0,102 | 9,913 | 0,501 |
| hsa-miR-761 | MIMAT0010364 | -0,994 | 0,246 | -1,010 | 0,424 | 1,568 | 0,048 |
| hsa-miR-888-3p | MIMAT0004917 | 1,197 | 0,508 | 1,514 | 0,195 | ND | - |
| hsa-miR-892b | MIMAT0004918 | 1,084 | 0,319 | 1,015 | 0,460 | ND | - |

**Supplementary Table 3.** List of 180 genes targeted by 41 miRNAs that modulate APP metabolism (see Supplementary Table 2 and (Chapuis et al. 2017) for the list of miRNAs). Sofware predicted the same targeted genes are also reported.

| miRNA | MIMAT | Gene_ID | Gene_name | Soft_DB |
|---|---|---|---|---|
| hsa-miR-1343-3p | MIMAT0019776 | 89796 | NAV1 | microt_cds,mirdb,mirtarbase,targetscan |
| hsa-miR-16-1-3p | MIMAT0004489 | 2247 | FGF2 | miranda,rna22,targetminer,targetscan |
| | | 3895 | KTN1 | microt_cds,miranda,mirdb,targetscan |
| | | 5935 | RBM3 | microt_cds,miranda,mirdb,targetscan |
| | | 9662 | CEP135 | microt_cds,miranda,mirdb,rna22,targetscan |
| | | 10137 | RBM12 | microt_cds,miranda,targetminer,targetscan |
| | | 10352 | WARS2 | microt_cds,miranda,targetminer,targetscan |
| | | 51496 | CTDSPL2 | miranda,mirdb,targetminer,targetscan |
| | | 51719 | CAB39 | microt_cds,miranda,mirdb,targetminer,targetscan |
| | | 55608 | ANKRD10 | miranda,rna22,targetminer,targetscan |
| hsa-miR-181b-3p | MIMAT0022692 | 867 | CBL | microt_cds,mirdb,targetminer,targetscan |
| | | 2908 | NR3C1 | microt_cds,mirdb,rna22,targetminer,targetscan |
| | | 23291 | FBXW11 | microt_cds,mirdb,targetminer,targetscan |
| | | 26054 | SENP6 | microt_cds,mirdb,targetminer,targetscan |
| hsa-miR-190a-3p | MIMAT0026482 | 1316 | KLF6 | microt_cds,mirdb,mirtarbase,targetscan |
| | | 2908 | NR3C1 | microt_cds,mirdb,mirtarbase,targetscan |
| | | 5504 | PPP1R2 | microt_cds,mirdb,mirtarbase,targetscan |
| | | 6801 | STRN | microt_cds,mirdb,mirtarbase,targetscan |
| | | 7994 | KAT6A | microt_cds,mirdb,mirtarbase,targetscan |
| | | 8453 | CUL2 | microt_cds,mirdb,mirtarbase,targetscan |
| | | 51454 | GULP1 | microt_cds,mirdb,mirtarbase,targetscan |
| | | 54764 | ZRANB1 | microt_cds,mirdb,mirtarbase,targetscan |
| | | 57551 | TAOK1 | microt_cds,mirdb,mirtarbase,targetscan |
| | | 129684 | CNTNAP5 | microt_cds,mirdb,mirtarbase,targetscan |
| hsa-miR-193a-3p | MIMAT0000459 | 2843 | GPR20 | microt_cds,miranda,mirdb,targetscan |
| | | 6477 | SIAH1 | microt_cds,miranda,mirdb,rna22,targetscan |
| | | 10220 | GDF11 | microt_cds,miranda,mirdb,mirtarbase,targetscan |
| | | 51496 | CTDSPL2 | microt_cds,miranda,mirdb,targetminer,targetscan |
| | | 51719 | CAB39 | miranda,mirdb,targetminer,targetscan |
| | | 65108 | MARCKSL1 | microt_cds,miranda,mirdb,targetscan |
| hsa-miR-193b-3p | MIMAT0002819 | 2843 | GPR20 | microt_cds,miranda,mirdb,targetscan |
| | | 6477 | SIAH1 | microt_cds,miranda,mirdb,targetscan |
| | | 10220 | GDF11 | microt_cds,miranda,mirdb,mirtarbase,targetscan |
| | | 51496 | CTDSPL2 | microt_cds,miranda,mirdb,mirtarbase,targetminer,target scan |
| | | 51719 | CAB39 | miranda,mirdb,targetminer,targetscan |
| | | 57190 | SELENON | miranda,mirdb,mirtarbase,targetscan |
| | | 65108 | MARCKSL1 | microt_cds,miranda,mirdb,targetscan |
| hsa-miR-194-3p | MIMAT0004671 | 5829 | PXN | miranda,rna22,targetminer,targetscan |
| | | 10057 | ABCC5 | miranda,rna22,targetminer,targetscan |
| | | 10439 | OLFM1 | miranda,mirdb,targetminer,targetscan |

| | | | | |
|---|---|---|---|---|
| | | 79183 | TTPAL | microt_cds,mirdb,rna22,targetscan |
| | | 89796 | NAV1 | miranda,rna22,targetminer,targetscan |
| | | 90161 | HS6ST2 | miranda,rna22,targetminer,targetscan |
| | | 91452 | ACBD5 | miranda,mirdb,targetminer,targetscan |
| | | 143384 | CACUL1 | microt_cds,miranda,rna22,targetminer,targetscan |
| | | 146691 | TOM1L2 | mirdb,rna22,targetminer,targetscan |
| hsa-miR-200b-3p | MIMAT0000318 | 3930 | LBR | microt_cds,miranda,mirdb,targetscan |
| | | 4884 | NPTX1 | microt_cds,miranda,mirdb,targetscan |
| | | 4908 | NTF3 | microt_cds,miranda,mirdb,targetscan |
| | | 6405 | SEMA3F | microt_cds,miranda,mirdb,targetscan |
| | | 6477 | SIAH1 | microt_cds,miranda,mirdb,targetscan |
| | | 7093 | TLL2 | microt_cds,miranda,targetminer,targetscan |
| | | 7532 | YWHAG | microt_cds,miranda,mirdb,targetminer,targetscan |
| | | 10777 | ARPP21 | microt_cds,miranda,mirdb,rna22,targetscan |
| | | 10979 | FERMT2 | microt_cds,miranda,mirtarbase,targetscan |
| | | 11183 | MAP4K5 | microt_cds,miranda,mirdb,targetminer,targetscan |
| | | 22841 | RAB11FIP2 | microt_cds,miranda,mirdb,targetminer,targetscan |
| | | 23328 | SASH1 | microt_cds,miranda,mirdb,targetscan |
| | | 23362 | PSD3 | mirdb,mirtarbase,targetminer,targetscan |
| | | 51496 | CTDSPL2 | microt_cds,miranda,mirdb,targetscan |
| | | 51719 | CAB39 | miranda,mirdb,mirtarbase,targetscan |
| | | 57826 | RAP2C | microt_cds,miranda,mirdb,targetscan |
| | | 284695 | ZNF326 | microt_cds,miranda,mirdb,targetscan |
| hsa-miR-2115-5p | MIMAT0011158 | 2196 | FAT2 | miranda,rna22,targetminer,targetscan |
| | | 9228 | DLGAP2 | miranda,mirdb,targetminer,targetscan |
| | | 11108 | PRDM4 | microt_cds,miranda,rna22,targetscan |
| | | 22992 | KDM2A | microt_cds,miranda,rna22,targetscan |
| | | 51060 | TXNDC12 | microt_cds,miranda,mirdb,targetscan |
| | | 89797 | NAV2 | microt_cds,miranda,mirdb,targetscan |
| | | 117177 | RAB3IP | miranda,mirtarbase,targetminer,targetscan |
| | | 219899 | TBCEL | microt_cds,miranda,mirdb,targetscan |
| hsa-miR-221-3p | MIMAT0000278 | 405 | ARNT | microt_cds,mirdb,mirtarbase,rna22,targetminer,targetscan |
| | | 4908 | NTF3 | microt_cds,miranda,mirdb,targetminer,targetscan |
| | | 6096 | RORB | microt_cds,miranda,targetminer,targetscan |
| | | 7532 | YWHAG | microt_cds,miranda,mirdb,rna22,targetscan |
| | | 9019 | MPZL1 | microt_cds,miranda,rna22,targetscan |
| | | 10979 | FERMT2 | microt_cds,miranda,mirdb,targetscan |
| | | 25874 | MPC2 | microt_cds,miranda,rna22,targetscan |
| | | 360023 | ZBTB41 | miranda,mirdb,rna22,targetscan |
| hsa-miR-222-3p | MIMAT0000279 | 405 | ARNT | microt_cds,mirdb,rna22,targetminer,targetscan |
| | | 4908 | NTF3 | microt_cds,miranda,mirdb,targetminer,targetscan |
| | | 6096 | RORB | microt_cds,miranda,targetminer,targetscan |
| | | 7532 | YWHAG | microt_cds,miranda,mirdb,mirtarbase,rna22,targetscan |
| | | 9019 | MPZL1 | microt_cds,miranda,rna22,targetscan |

| | | | | |
|---|---|---|---|---|
| | | 10979 | FERMT2 | microt_cds,miranda,mirdb,targetscan |
| | | 56977 | STOX2 | miranda,mirtarbase,rna22,targetscan |
| hsa-miR-3160-3p | MIMAT0015034 | 3213 | HOXB3 | microt_cds,miranda,rna22,targetscan |
| | | 10777 | ARPP21 | miranda,mirdb,targetminer,targetscan |
| | | 23087 | TRIM35 | miranda,mirdb,rna22,targetscan |
| | | 23299 | BICD2 | mirtarbase,rna22,targetminer,targetscan |
| | | 26468 | LHX6 | miranda,rna22,targetminer,targetscan |
| | | 55219 | TMEM57 | miranda,mirdb,targetminer,targetscan |
| hsa-miR-3160-5p | MIMAT0019212 | 5087 | PBX1 | microt_cds,mirdb,rna22,targetscan |
| | | 23299 | BICD2 | microt_cds,rna22,targetminer,targetscan |
| | | 23328 | SASH1 | microt_cds,mirdb,rna22,targetscan |
| hsa-miR-3179 | MIMAT0015056 | 3202 | HOXA5 | microt_cds,miranda,targetminer,targetscan |
| | | 3219 | HOXB9 | microt_cds,miranda,mirdb,targetminer,targetscan |
| | | 5087 | PBX1 | miranda,mirdb,targetminer,targetscan |
| | | 6405 | SEMA3F | miranda,mirtarbase,rna22,targetminer,targetscan |
| | | 6844 | VAMP2 | microt_cds,miranda,mirdb,targetminer,targetscan |
| | | 10509 | SEMA4B | microt_cds,miranda,targetminer,targetscan |
| | | 22992 | KDM2A | microt_cds,miranda,targetminer,targetscan |
| | | 57523 | NYNRIN | microt_cds,miranda,mirdb,rna22,targetscan |
| | | 64855 | FAM129B | miranda,mirtarbase,targetminer,targetscan |
| | | 80320 | SP6 | microt_cds,miranda,mirdb,targetminer,targetscan |
| | | 81566 | CSRNP2 | microt_cds,miranda,mirdb,targetminer,targetscan |
| | | 81605 | URM1 | microt_cds,miranda,mirdb,mirtarbase,targetminer,target scan |
| | | 89797 | NAV2 | miranda,mirtarbase,targetminer,targetscan |
| | | 92126 | DSEL | miranda,mirdb,targetminer,targetscan |
| | | 286046 | XKR6 | microt_cds,miranda,mirdb,rna22,targetminer,targetscan |
| hsa-miR-3188 | MIMAT0015070 | 215 | ABCD1 | microt_cds,miranda,mirdb,rna22 |
| | | 1910 | EDNRB | microt_cds,miranda,targetminer,targetscan |
| | | 5830 | PEX5 | microt_cds,miranda,mirdb,targetminer,targetscan |
| | | 7994 | KAT6A | microt_cds,miranda,rna22,targetminer,targetscan |
| | | 8994 | LIMD1 | microt_cds,rna22,targetminer,targetscan |
| | | 11261 | CHP1 | microt_cds,mirdb,rna22,targetminer,targetscan |
| | | 25963 | TMEM87A | miranda,rna22,targetminer,targetscan |
| | | 51804 | SIX4 | miranda,mirdb,rna22,targetminer,targetscan |
| | | 56111 | PCDHGA4 | microt_cds,miranda,mirdb,targetminer,targetscan |
| | | 56112 | PCDHGA3 | microt_cds,miranda,mirdb,rna22,targetminer,targetscan |
| | | 56255 | TMX4 | microt_cds,miranda,mirdb,targetminer,targetscan |
| | | 57337 | SENP7 | microt_cds,miranda,mirdb,rna22,targetminer,targetscan |
| | | 57551 | TAOK1 | microt_cds,miranda,rna22,targetminer,targetscan |
| | | 79183 | TTPAL | mirdb,rna22,targetminer,targetscan |
| | | 79668 | PARP8 | microt_cds,rna22,targetminer,targetscan |
| | | 399664 | MEX3D | microt_cds,miranda,rna22,targetscan |
| hsa-miR-32-3p | MIMAT0004505 | 36 | ACADSB | microt_cds,miranda,targetminer,targetscan |
| | | 6648 | SOD2 | microt_cds,miranda,rna22,targetscan |

| | | | | |
|---|---|---|---|---|
| | | 7994 | KAT6A | microt_cds,miranda,targetminer,targetscan |
| | | 10371 | SEMA3A | microt_cds,miranda,mirdb,targetminer,targetscan |
| | | 23291 | FBXW11 | microt_cds,miranda,mirdb,targetminer,targetscan |
| | | 26054 | SENP6 | microt_cds,miranda,targetminer,targetscan |
| | | 51454 | GULP1 | microt_cds,miranda,targetminer,targetscan |
| | | 51804 | SIX4 | microt_cds,miranda,targetminer,targetscan |
| | | 55432 | YOD1 | microt_cds,miranda,mirdb,targetminer,targetscan |
| | | 57337 | SENP7 | microt_cds,miranda,mirdb,targetminer,targetscan |
| | | 66008 | TRAK2 | microt_cds,miranda,targetminer,targetscan |
| | | 79183 | TTPAL | miranda,mirdb,targetminer,targetscan |
| | | 81034 | SLC25A32 | miranda,rna22,targetminer,targetscan |
| | | 90161 | HS6ST2 | microt_cds,miranda,targetminer,targetscan |
| | | 91452 | ACBD5 | microt_cds,miranda,mirdb,targetminer,targetscan |
| | | 116931 | MED12L | microt_cds,miranda,mirtarbase,targetscan |
| | | 117178 | SSX2IP | microt_cds,miranda,targetminer,targetscan |
| | | 137970 | UNC5D | microt_cds,miranda,targetminer,targetscan |
| | | 346007 | EYS | microt_cds,miranda,mirdb,targetscan |
| hsa-miR-3529-5p | MIMAT0019828 | 10371 | SEMA3A | mirdb,rna22,targetminer,targetscan |
| | | 80267 | EDEM3 | mirdb,mirtarbase,rna22,targetminer,targetscan |
| | | 285636 | C5orf51 | microt_cds,mirtarbase,targetminer,targetscan |
| hsa-miR-3616-5p | MIMAT0017995 | 23362 | PSD3 | microt_cds,mirdb,targetminer,targetscan |
| | | 92126 | DSEL | microt_cds,mirdb,targetminer,targetscan |
| | | 339745 | SPOPL | microt_cds,mirdb,targetminer,targetscan |
| hsa-miR-3663-3p | MIMAT0018085 | 4826 | NNAT | microt_cds,mirdb,targetminer,targetscan |
| | | 51496 | CTDSPL2 | microt_cds,mirdb,targetminer,targetscan |
| hsa-miR-3668 | MIMAT0018091 | 577 | ADGRB3 | microt_cds,mirdb,targetminer,targetscan |
| | | 8453 | CUL2 | microt_cds,mirdb,targetminer,targetscan |
| | | 10971 | YWHAQ | microt_cds,mirdb,targetminer,targetscan |
| | | 25963 | TMEM87A | microt_cds,mirdb,targetminer,targetscan |
| hsa-miR-3977 | MIMAT0019362 | 22841 | RAB11FIP2 | microt_cds,mirdb,targetminer,targetscan |
| hsa-miR-4429 | MIMAT0018944 | 2045 | EPHA7 | microt_cds,rna22,targetminer,targetscan |
| | | 3664 | IRF6 | microt_cds,mirdb,targetminer,targetscan |
| | | 9053 | MAP7 | microt_cds,mirdb,targetminer,targetscan |
| | | 22841 | RAB11FIP2 | microt_cds,mirdb,targetminer,targetscan |
| | | 23328 | SASH1 | microt_cds,mirdb,targetminer,targetscan |
| | | 26036 | ZNF451 | mirtarbase,rna22,targetminer,targetscan |
| | | 80014 | WWC2 | microt_cds,mirdb,targetminer,targetscan |
| | | 117177 | RAB3IP | microt_cds,rna22,targetminer,targetscan |
| | | 339745 | SPOPL | microt_cds,mirdb,targetminer,targetscan |
| | | 388531 | RGS9BP | microt_cds,mirdb,targetminer,targetscan |
| hsa-miR-4477a | MIMAT0019004 | 23291 | FBXW11 | microt_cds,mirdb,targetminer,targetscan |
| | | 80267 | EDEM3 | microt_cds,mirdb,targetminer,targetscan |
| | | 399664 | MEX3D | microt_cds,mirdb,targetminer,targetscan |
| hsa-miR-4773 | MIMAT0019928 | 2334 | AFF2 | microt_cds,mirtarbase,targetminer,targetscan |
| | | 3358 | HTR2C | microt_cds,mirdb,rna22,targetminer,targetscan |

| | | | | |
|---|---|---|---|---|
| | | 4152 | MBD1 | microt_cds,mirdb,targetminer,targetscan |
| | | 9880 | ZBTB39 | microt_cds,mirdb,mirtarbase,targetminer,targetscan |
| | | 10180 | RBM6 | microt_cds,mirdb,targetminer,targetscan |
| | | 23328 | SASH1 | microt_cds,mirdb,mirtarbase,rna22,targetminer,targetscan |
| hsa-miR-488-5p | MIMAT0002804 | 36 | ACADSB | miranda,mirdb,targetminer,targetscan |
| | | 1910 | EDNRB | microt_cds,miranda,targetminer,targetscan |
| | | 2908 | NR3C1 | miranda,rna22,targetminer,targetscan |
| | | 3597 | IL13RA1 | microt_cds,miranda,mirdb,targetminer,targetscan |
| | | 10564 | ARFGEF2 | microt_cds,miranda,mirdb,targetminer,targetscan |
| hsa-miR-506-3p | MIMAT0002878 | 867 | CBL | microt_cds,mirdb,targetminer,targetscan |
| | | 1316 | KLF6 | microt_cds,miranda,mirdb,targetminer,targetscan |
| | | 1910 | EDNRB | microt_cds,miranda,mirdb,targetscan |
| | | 2029 | ENSA | miranda,rna22,targetminer,targetscan |
| | | 2892 | GRIA3 | microt_cds,miranda,mirdb,targetminer |
| | | 2908 | NR3C1 | microt_cds,miranda,targetminer,targetscan |
| | | 3688 | ITGB1 | microt_cds,miranda,mirdb,targetminer |
| | | 6648 | SOD2 | microt_cds,miranda,targetminer,targetscan |
| | | 6744 | SSFA2 | microt_cds,miranda,mirdb,targetscan |
| | | 8715 | NOL4 | microt_cds,miranda,targetminer,targetscan |
| | | 8994 | LIMD1 | microt_cds,mirdb,targetminer,targetscan |
| | | 9697 | TRAM2 | microt_cds,mirdb,targetminer,targetscan |
| | | 10564 | ARFGEF2 | microt_cds,miranda,mirdb,targetminer,targetscan |
| | | 10788 | IQGAP2 | miranda,mirdb,targetminer,targetscan |
| | | 11261 | CHP1 | microt_cds,mirdb,targetminer,targetscan |
| | | 51804 | SIX4 | microt_cds,miranda,mirdb,targetminer,targetscan |
| | | 58533 | SNX6 | microt_cds,miranda,targetminer,targetscan |
| | | 83871 | RAB34 | microt_cds,miranda,mirdb,targetminer,targetscan |
| | | 89796 | NAV1 | microt_cds,rna22,targetminer,targetscan |
| | | 137970 | UNC5D | microt_cds,miranda,targetminer,targetscan |
| | | 143384 | CACUL1 | microt_cds,mirdb,targetminer,targetscan |
| hsa-miR-511-3p | MIMAT0026606 | 10371 | SEMA3A | microt_cds,mirdb,rna22,targetscan |
| hsa-miR-513c-3p | MIMAT0022728 | 2908 | NR3C1 | microt_cds,mirdb,targetminer,targetscan |
| | | 7994 | KAT6A | microt_cds,mirdb,targetminer,targetscan |
| | | 10484 | SEC23A | microt_cds,mirdb,targetminer,targetscan |
| | | 64786 | TBC1D15 | microt_cds,mirdb,targetminer,targetscan |
| | | 127428 | TCEANC2 | microt_cds,mirdb,targetminer,targetscan |
| | | 137970 | UNC5D | microt_cds,mirdb,targetminer,targetscan |
| hsa-miR-5193 | MIMAT0021124 | 59271 | EVA1C | microt_cds,mirdb,rna22,targetscan |
| | | 89796 | NAV1 | microt_cds,mirdb,mirtarbase,rna22,targetminer,targetscan |
| hsa-miR-5579-5p | MIMAT0022269 | 10371 | SEMA3A | microt_cds,mirdb,targetminer,targetscan |
| | | 23429 | RYBP | microt_cds,mirdb,targetminer,targetscan |
| hsa-miR-5582-5p | MIMAT0022279 | 867 | CBL | microt_cds,mirdb,rna22,targetminer,targetscan |
| | | 2908 | NR3C1 | microt_cds,mirtarbase,targetminer,targetscan |
| | | 3688 | ITGB1 | microt_cds,mirdb,targetminer,targetscan |

| | | | | |
|---|---|---|---|---|
| | | 5504 | PPP1R2 | microt_cds,mirdb,targetminer,targetscan |
| | | 10788 | IQGAP2 | microt_cds,mirdb,targetminer,targetscan |
| | | 11261 | CHP1 | microt_cds,mirdb,targetminer,targetscan |
| | | 23291 | FBXW11 | microt_cds,mirdb,targetminer,targetscan |
| | | 51804 | SIX4 | microt_cds,mirdb,targetminer,targetscan |
| | | 57658 | CALCOCO1 | microt_cds,mirdb,targetminer,targetscan |
| | | 137970 | UNC5D | microt_cds,mirdb,targetminer,targetscan |
| | | 143384 | CACUL1 | microt_cds,mirdb,targetminer,targetscan |
| hsa-miR-582-5p | MIMAT0003247 | 2247 | FGF2 | microt_cds,miranda,mirtarbase,targetminer,targetscan |
| | | 10979 | FERMT2 | microt_cds,miranda,mirdb,targetminer,targetscan |
| | | 23299 | BICD2 | microt_cds,miranda,mirdb,targetminer,targetscan |
| | | 23362 | PSD3 | microt_cds,mirdb,targetminer,targetscan |
| | | 51060 | TXNDC12 | microt_cds,miranda,mirdb,targetminer,targetscan |
| | | 57826 | RAP2C | microt_cds,miranda,mirtarbase,targetminer,targetscan |
| | | 80790 | CMIP | microt_cds,miranda,mirdb,targetminer,targetscan |
| | | 81566 | CSRNP2 | microt_cds,miranda,targetminer,targetscan |
| | | 151742 | PPM1L | microt_cds,miranda,mirdb,targetscan |
| | | 222171 | PRR15 | microt_cds,miranda,mirdb,targetscan |
| | | 360023 | ZBTB41 | microt_cds,miranda,mirdb,targetminer,targetscan |
| hsa-miR-6783-3p | MIMAT0027467 | 867 | CBL | microt_cds,mirdb,rna22,targetscan |
| | | 84448 | ABLIM2 | microt_cds,mirdb,rna22,targetscan |
| | | 89796 | NAV1 | microt_cds,mirdb,mirtarbase,rna22,targetscan |
| | | 148254 | ZNF555 | microt_cds,mirdb,rna22,targetscan |
| | | 376267 | RAB15 | microt_cds,mirtarbase,rna22,targetscan |
| hsa-miR-6884-5p | MIMAT0027668 | 414 | ARSD | microt_cds,mirdb,rna22,targetscan |
| | | 11261 | CHP1 | microt_cds,mirdb,rna22,targetscan |
| | | 57551 | TAOK1 | microt_cds,mirdb,rna22,targetscan |
| | | 79668 | PARP8 | microt_cds,mirdb,rna22,targetscan |
| | | 89796 | NAV1 | microt_cds,mirdb,mirtarbase,rna22,targetscan |
| | | 90594 | ZNF439 | microt_cds,mirtarbase,rna22,targetscan |
| | | 129685 | TAF8 | microt_cds,mirtarbase,rna22,targetscan |
| | | 220074 | LRTOMT | microt_cds,mirdb,rna22,targetscan |
| | | 245972 | ATP6V0D2 | microt_cds,mirdb,mirtarbase,targetscan |
| hsa-miR-7-5p | MIMAT0000252 | 142 | PARP1 | microt_cds,miranda,mirdb,mirtarbase,targetscan |
| | | 867 | CBL | microt_cds,mirdb,rna22,targetminer,targetscan |
| | | 2029 | ENSA | miranda,mirtarbase,rna22,targetminer,targetscan |
| | | 2768 | GNA12 | microt_cds,miranda,rna22,targetminer,targetscan |
| | | 6500 | SKP1 | microt_cds,miranda,mirdb,targetminer,targetscan |
| | | 6536 | SLC6A9 | microt_cds,miranda,mirdb,mirtarbase,targetscan |
| | | 8994 | LIMD1 | microt_cds,miranda,mirdb,targetminer,targetscan |
| | | 9697 | TRAM2 | microt_cds,mirtarbase,targetminer,targetscan |
| | | 11261 | CHP1 | microt_cds,mirdb,targetminer,targetscan |
| | | 23211 | ZC3H4 | microt_cds,miranda,mirdb,mirtarbase,targetscan |
| | | 54107 | POLE3 | microt_cds,miranda,mirtarbase,targetscan |
| | | 55432 | YOD1 | miranda,mirtarbase,targetminer,targetscan |

| | | | | |
|---|---|---|---|---|
| | | 55591 | VEZT | microt_cds,miranda,rna22,targetscan |
| | | 56255 | TMX4 | microt_cds,miranda,targetminer,targetscan |
| | | 89796 | NAV1 | miranda,mirtarbase,targetminer,targetscan |
| | | 91452 | ACBD5 | microt_cds,rna22,targetminer,targetscan |
| | | 129685 | TAF8 | microt_cds,miranda,mirtarbase,rna22,targetscan |
| | | 134353 | LSM11 | microt_cds,miranda,targetminer,targetscan |
| hsa-miR-7151-3p | MIMAT0028213 | 54751 | FBLIM1 | microt_cds,mirdb,rna22,targetscan |
| hsa-miR-7160-3p | MIMAT0028231 | 84957 | RELT | microt_cds,mirdb,rna22,targetscan |
| hsa-miR-7160-5p | MIMAT0028230 | 4293 | MAP3K9 | microt_cds,mirdb,rna22,targetscan |
| | | 10350 | ABCA9 | microt_cds,mirdb,rna22,targetscan |
| | | 23299 | BICD2 | mirdb,mirtarbase,rna22,targetscan |
| | | 57002 | YAE1D1 | microt_cds,mirdb,rna22,targetscan |
| hsa-miR-761 | MIMAT0010364 | 867 | CBL | microt_cds,mirdb,targetminer,targetscan |
| | | 1106 | CHD2 | miranda,mirdb,targetminer,targetscan |
| | | 2909 | ARHGAP35 | microt_cds,mirdb,rna22,targetminer,targetscan |
| | | 5076 | PAX2 | microt_cds,rna22,targetminer,targetscan |
| | | 5332 | PLCB4 | miranda,mirdb,targetminer,targetscan |
| | | 5338 | PLD2 | microt_cds,miranda,mirdb,targetminer,targetscan |
| | | 9543 | IGDCC3 | miranda,rna22,targetminer,targetscan |
| | | 9832 | JAKMIP2 | microt_cds,miranda,mirdb,targetminer,targetscan |
| | | 10057 | ABCC5 | microt_cds,miranda,mirdb,targetminer |
| | | 10564 | ARFGEF2 | microt_cds,miranda,targetminer,targetscan |
| | | 11261 | CHP1 | microt_cds,rna22,targetminer,targetscan |
| | | 23035 | PHLPP2 | mirtarbase,rna22,targetminer,targetscan |
| | | 23429 | RYBP | microt_cds,miranda,targetminer,targetscan |
| | | 25963 | TMEM87A | microt_cds,miranda,targetminer,targetscan |
| | | 27244 | SESN1 | microt_cds,miranda,mirdb,rna22,targetminer,targetscan |
| | | 55591 | VEZT | miranda,mirdb,targetminer,targetscan |
| | | 57614 | KIAA1468 | miranda,rna22,targetminer,targetscan |
| | | 64786 | TBC1D15 | miranda,rna22,targetminer,targetscan |
| | | 80218 | NAA50 | miranda,mirdb,targetminer,targetscan |
| | | 89796 | NAV1 | microt_cds,rna22,targetminer,targetscan |
| | | 117178 | SSX2IP | miranda,mirdb,targetminer,targetscan |
| | | 134353 | LSM11 | microt_cds,miranda,targetminer,targetscan |
| | | 143384 | CACUL1 | microt_cds,rna22,targetminer,targetscan |
| | | 196527 | ANO6 | miranda,mirdb,rna22,targetscan |
| | | 376267 | RAB15 | microt_cds,miranda,rna22,targetminer,targetscan |
| hsa-miR-888-3p | MIMAT0004917 | 3093 | UBE2K | miranda,mirdb,targetminer,targetscan |
| | | 3202 | HOXA5 | microt_cds,miranda,mirdb,targetscan |
| | | 7270 | TTF1 | microt_cds,miranda,mirdb,targetscan |
| | | 7544 | ZFY | miranda,mirdb,targetminer,targetscan |
| hsa-miR-892b | MIMAT0004918 | 7544 | ZFY | microt_cds,miranda,targetminer,targetscan |
| | | 7782 | SLC30A4 | microt_cds,miranda,mirdb,rna22,targetscan |
| | | 11183 | MAP4K5 | miranda,mirdb,targetminer,targetscan |
| | | 23299 | BICD2 | microt_cds,miranda,targetminer,targetscan |

| | | | | |
|---|---|---|---|---|
| | | 51496 | CTDSPL2 | microt_cds,miranda,targetminer,targetscan |
| | | 51704 | GPRC5B | miranda,mirdb,rna22,targetscan |
| | | 51780 | KDM3B | microt_cds,miranda,rna22,targetscan |

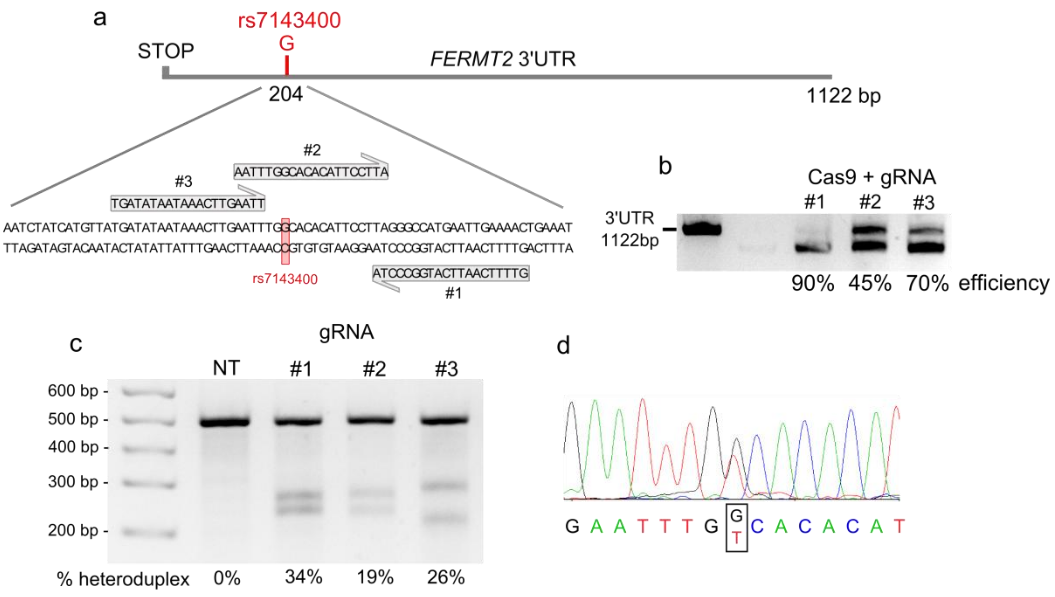


e

| Sequence | PAM | Score | Gene | Chromosome | Strand | Position | Mismatches |
|---|---|---|---|---|---|---|---|
| GTTTTCAATTCATGGCCCTA | AGG | 100.00 | *FERMT2* | chr14 | 1 | 52858159 | 0 |
| GCTTTCAGTTCATGGCCCTG | AAG | 1.87 | | chr2 | -1 | 2339188 | 3 |
| GTTTTCACTGCGTGGCCCTA | AGG | 1.13 | | chr1 | -1 | 224734940 | 3 |

**Supplementary Fig. 1. CRISPR/Cas9 editing of the HEK293 cell line for rs7143400**

**a.** Localization of the predicted gRNAs in the vicinity of the rs7143400. **b.** Assessment of gRNA + recombinant Cas9 complex efficiency for the cleavage using *FERMT2* 3’UTR cDNA. **c.** Assessment of gRNA + Cas9 complex efficiency in HEK293 using T7 endonuclease-based heteroduplex cleavage assay. HEK293 were transfected for 48 h with plasmids allowing Cas9 and gRNA (#1, #2 or #3) expression. **d.** Sanger sequencing showing the edition of a HEK293 clone for the rs7143400. **e.** The list of the potential off-targets predicted for the gRNA#1 (Score > 1 and maximum 3 mismatches). The integrity of the *FERMT2* 3’UTR sequence and the potential off-target sites were validated by sequencing.

a

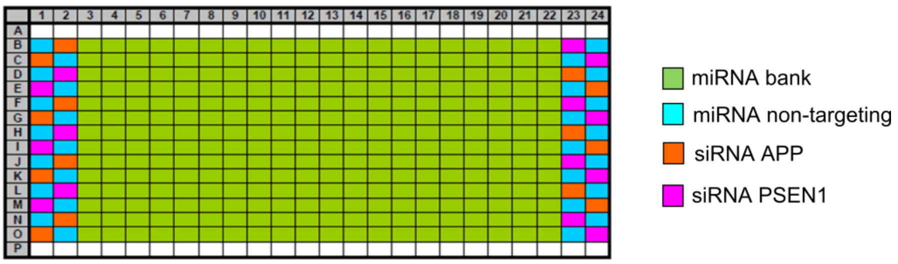


b

$$\beta = \frac{\mu_{PSEN1} - \mu_{APP}}{\sqrt{\sigma^2_{PSEN1} + \sigma^2_{APP}}}$$

Strictly standardized mean difference (SSMD, denoted as β)

c

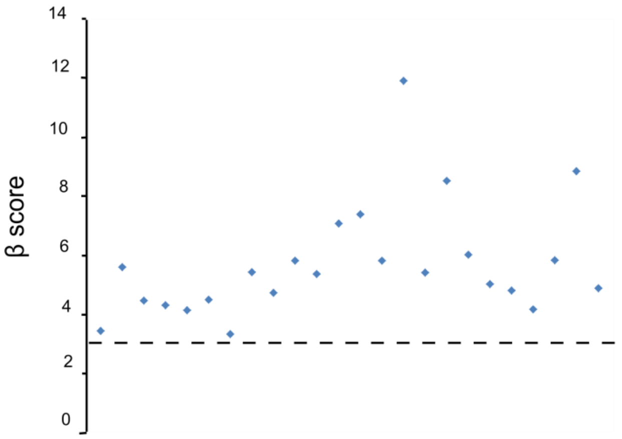


**Supplementary Fig. 2. Quality control for the high-content miRNA screening**

**a.** Well positions of miRNA and siRNA controls and the miRNA bank on the 384-well plate. The layouts were adjusted to systematically decrease the edge bias by alternating the spatial position of the controls so that they appear equally on each row and each available column. **b.** Formula used for to calculate the SSMD, which is a measure of the strength of the difference between two controls (PSEN1 and APP). μ and σ are the mean and standard deviation of the fluorescence intensity respectively. **c.** Graph showing the distribution of β scores for each plate. Dashed line indicates the quality threshold (β = 3).

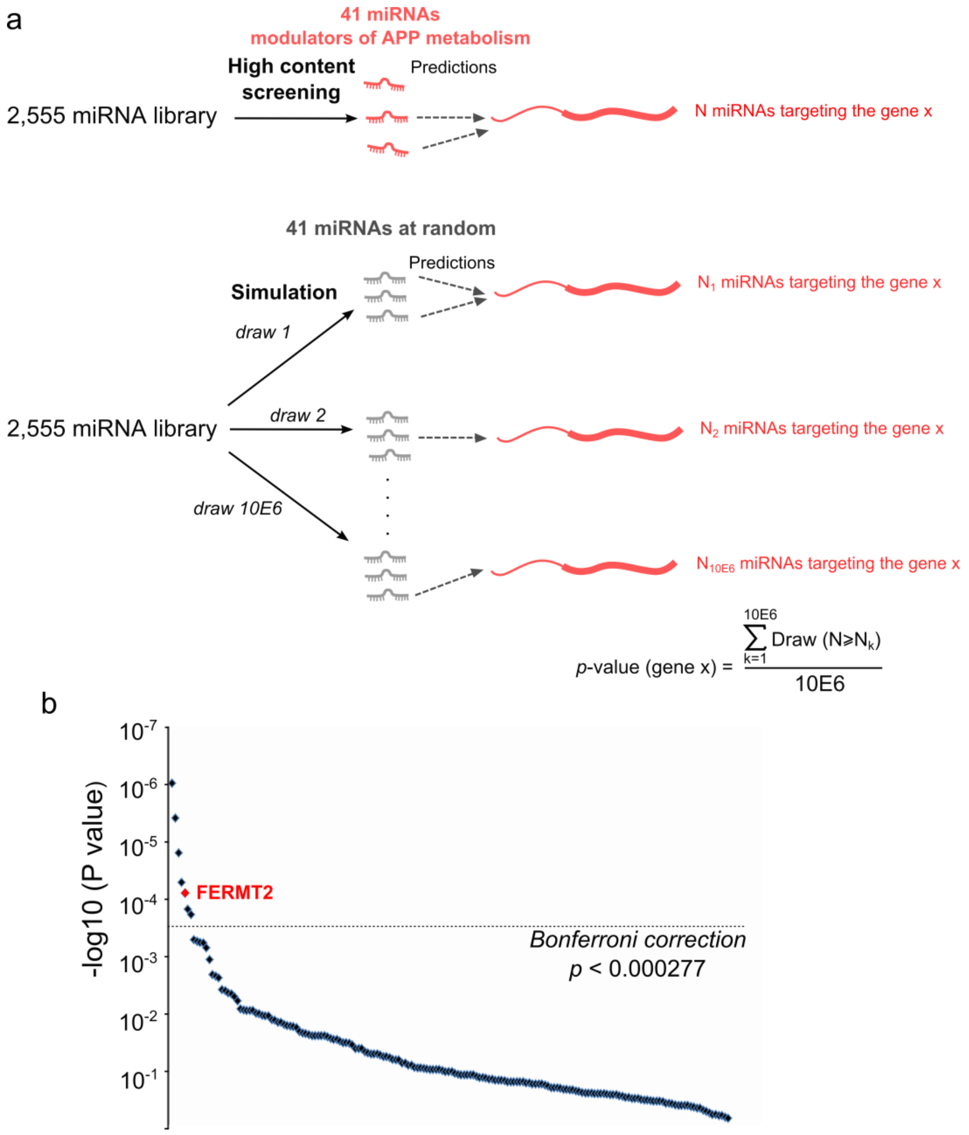


**Supplementary Fig. 3. Workflow to predict miRNA-targeted genes**

**a.** A graphical representation of the workflow used to attribute a *p*-value for each gene targeted by the best 41 miRNAs that are also able to modulate APP metabolism according to our genome-wide high-content siRNA screening (also see **Supplementary Table 2**). **b**. *p*-value distribution for the 180 genes targeted by the 41 miRNAs identified by HCS (also see **Supplementary Table 3**). In red, genes significantly enriched to be targeted by these 41 miRNAs after Bonferroni correction ($0.05 / 180 = 2.77\times10^{-4}$) including *FERMT2*, a genetic risk factor of AD.

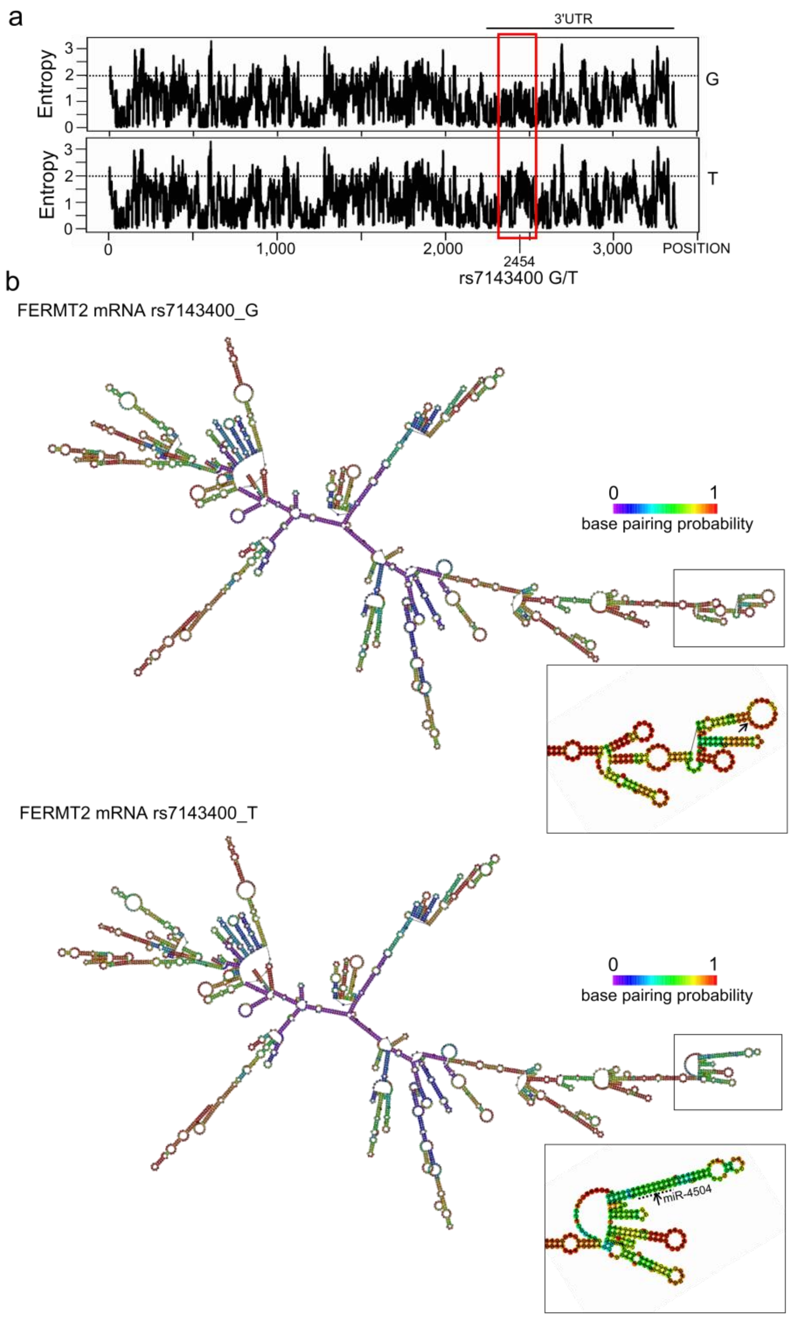


**Supplementary Fig. 4. Prediction of the impact of the rs7143400 allele on FERMT2 mRNA structure**

**a.** Comparison of the predicted entropy of FERMT2 mRNA structure according to the rs7143400 allele using RNAfold web server (http://rna.tbi.univie.ac.at/cgi-bin/RNAWebSuite/RNAfold.cgi). The red box notes the increase of entropy within the 3'UTR in the vicinity of the rs7143400. **b.** Impact of the rs7143400 on the predicted FERMT2 mRNA structure. Zoomed areas show the modified structure of rs7143400_T relative to rs7143400_G, including the sequence targeted by miR-4504.

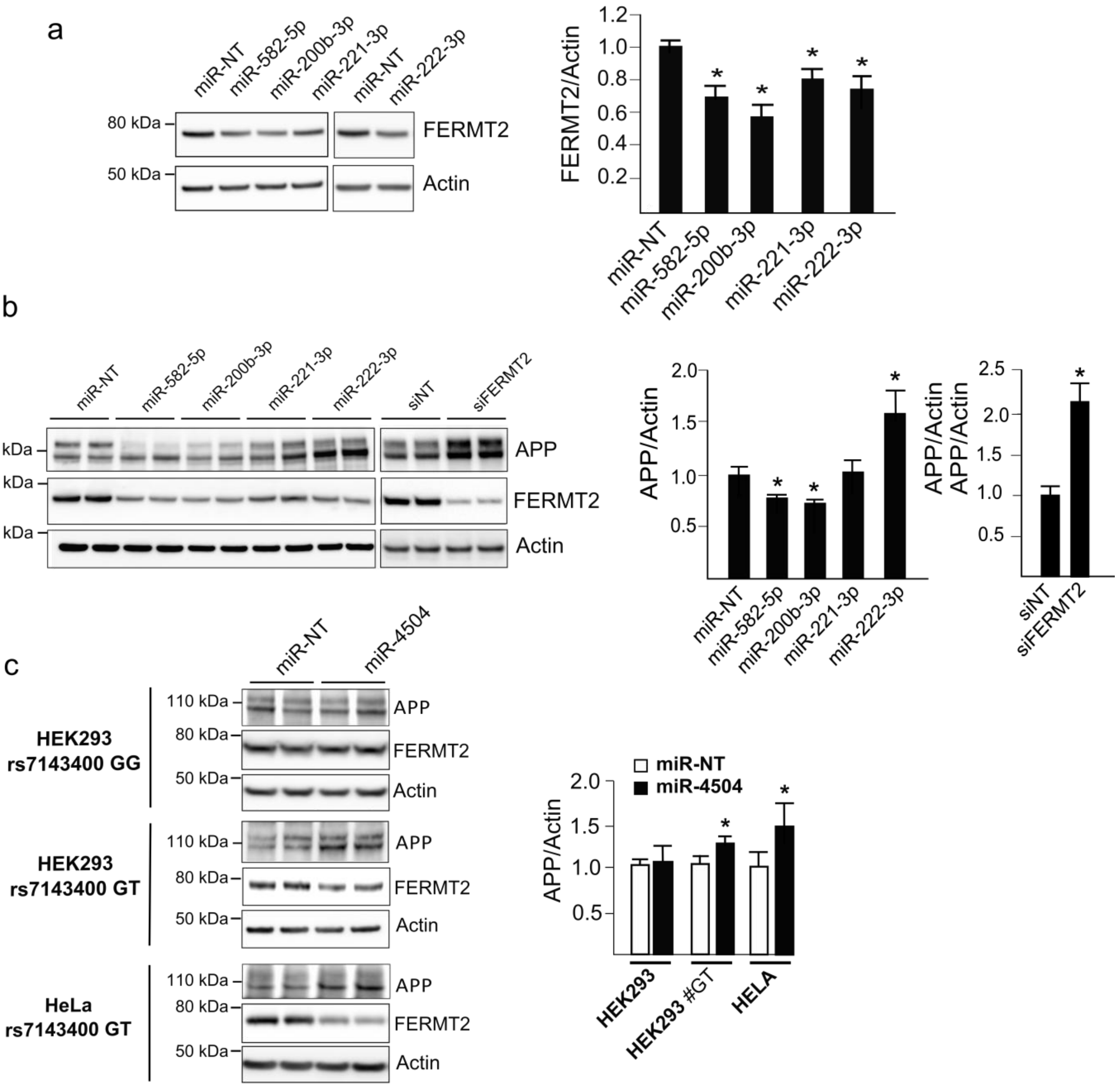


**Supplementary Fig. 5. Validation of the impact of miRNAs on endogenous FERMT2 and APP expression levels.**

**a.** Impact of miR-582-5p, miR-200b-3p, miR-221-3p and miR-222-3p on endogenous FERMT2 levels in hippocampi primary neuronal culture. FERMT2 expression levels were assessed by Western blot 72h after miRNAs transfection. Of note, such analysis was not possible for miR-4504, since the sequence encompassing rs7143400 is not present in the rodent. **b.** Impact of miR-582-5p, miR-200b-3p, miR-221-3p and miR-222-3p on endogenous FERMT2 and APP levels in HEK293 cell line. **c,** Endogenous FERMT2 and APP expression levels were assessed by Western blot using cell lines carrying the rs7143400-T allele (or not) following transient transfection with a non-targeting miR (miR-NT) or miR-4504. Bar charts show mean ± SD. Mann–Whitney test; * $p < 0.05$.

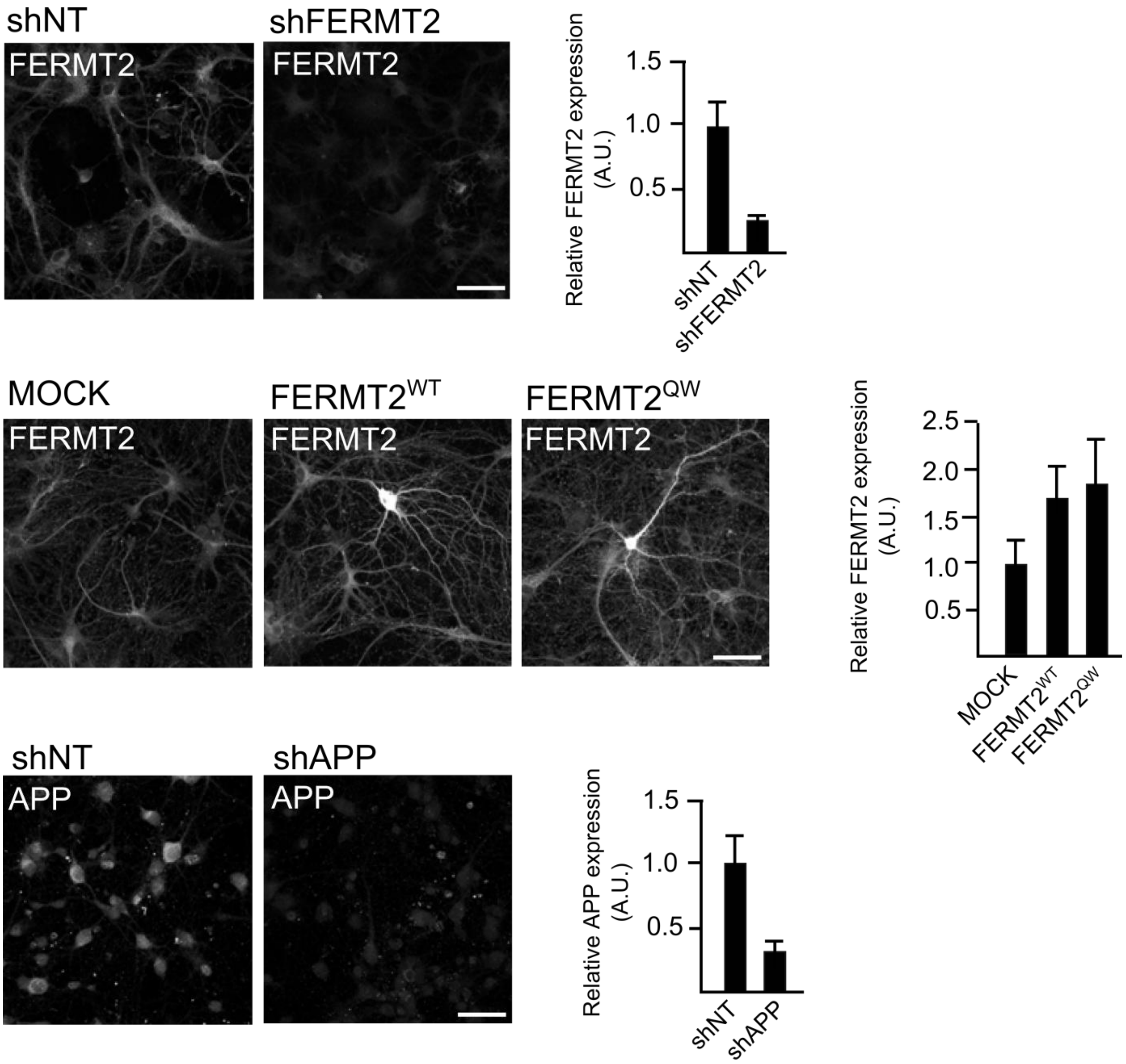


**Supplementary Fig. 6. Impact of lentiviral transduction on FERMT2 and APP expression levels.**

After transduction at DIV1 with the indicated lentivirus, expression levels of FERMT2 and APP were assessed by immunofluorescence using anti-FERMT2 or anti-APP antibodies, respectively. Scale bar= 10 µm. Bar charts show mean ± SD.

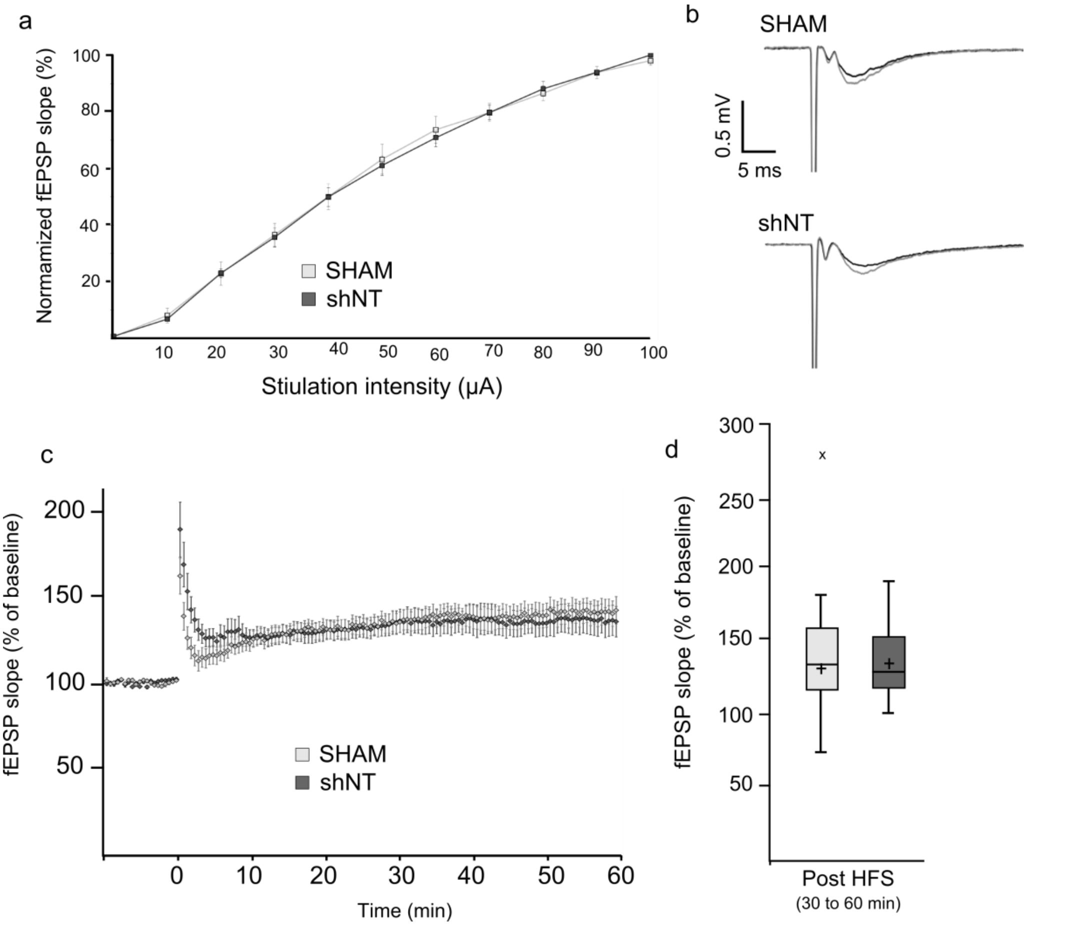


**Supplementary Fig. 7. Lentivirus injection does not alter electrophysiology recordings**

**a.** Normalized average slope of fEPSP evoked in hippocampal slices from animals injected with the indicated lentivirus or surgery procedures group (SHAM). N=5 animals/10 slices. **b**. Examples of recorded fEPSPs during baseline (black) and post LTP induction 40-60 min (grey). **c.** Time course of the average slope of elicited field responses following LTP induction by a tetanic stimulation protocol at hippocampal CA1 synapses from 5 mice after viral injection. Time-point 0 represents the delivery of tetanic stimulation. Slopes of each fEPSP were normalized to baseline and plotted against time. **d.** Box plots of the average slope response during 30-60 min post LTP induction. N = 5 mice; 2 slices per animal. × indicates an outlier.

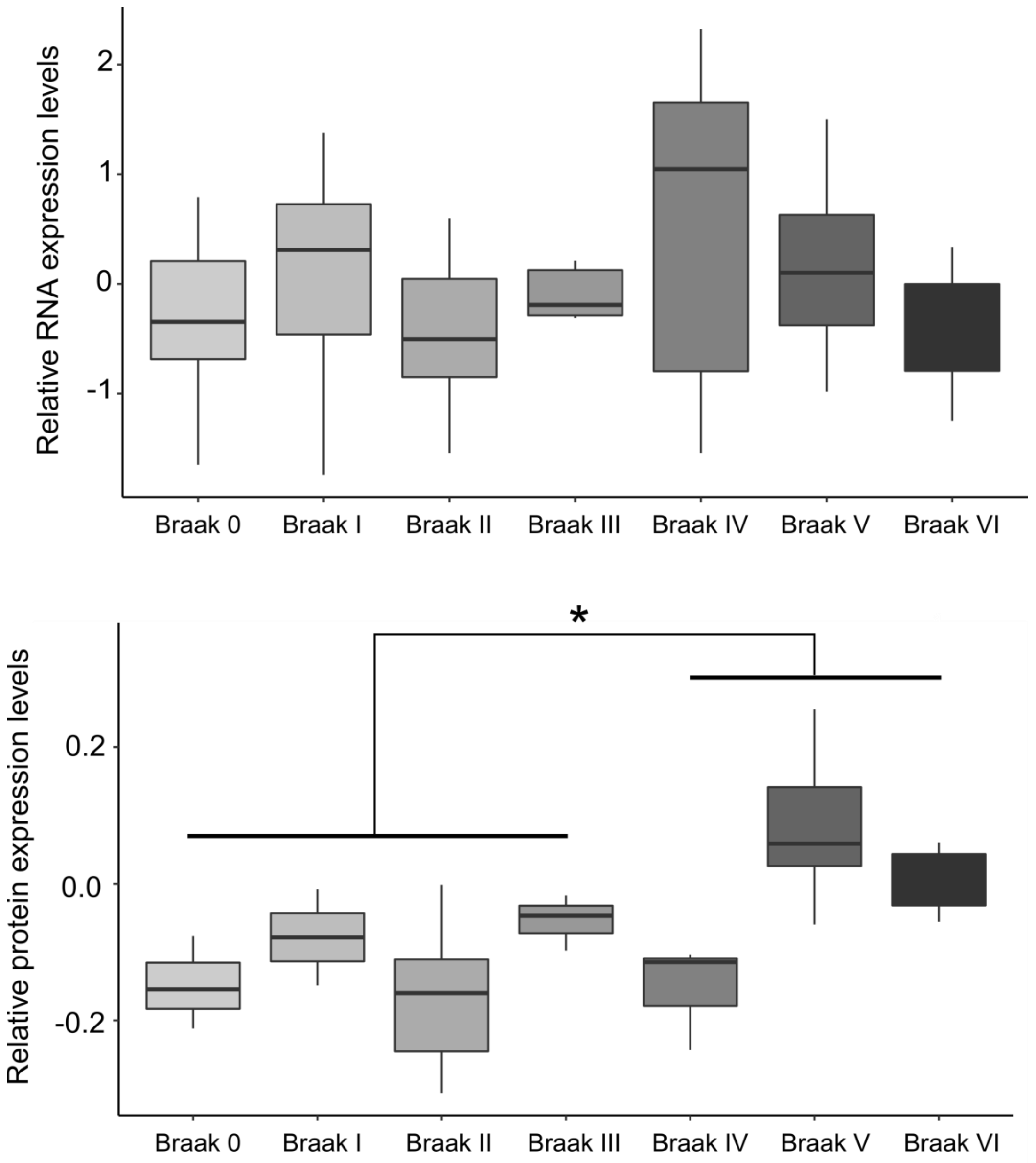


**Supplementary Fig. 8. Quantification of FERMT2 expression at the mRNA and protein levels in human brain samples**

FERMT2 protein level is significantly increased in post-mortem brains from patients at higher Braak stages. Statistics for global distribution, Kruskal-Wallis ANOVA (*p*-value = 0.0022), followed by Mann-Whitney U test, * *p*-value = 0.0008.
